# Systematic benchmarking of deep-learning methods for tertiary RNA structure prediction

**DOI:** 10.1101/2024.02.08.579037

**Authors:** Akash Bahai, Chee Keong Kwoh, Yuguang Mu, Yinghui Li

## Abstract

The 3D structure of RNA critically influences its functionality, and understanding this structure is vital for deciphering RNA biology. Experimental methods for determining RNA structures are labour-intensive, expensive, and time-consuming. Computational approaches have emerged as valuable tools, leveraging physics-based-principles and machine learning to predict RNA structures rapidly. Despite advancements, the accuracy of computational methods remains modest, especially when compared to protein structure prediction. Deep learning methods, while successful in protein structure prediction, have shown some promise for RNA structure prediction as well but face unique challenges. This study systematically benchmarks state-of-the-art deep learning methods for RNA structure prediction across diverse datasets. Our aim is to identify factors influencing performance variation, such as RNA family diversity, sequence length, RNA type, multiple sequence alignment (MSA) quality, and deep learning model architecture. We show that generally ML-based methods perform much better than non-ML methods on most RNA targets, although the performance difference isn’t substantial when working with unseen novel or synthetic RNAs. The quality of the MSA and secondary structure prediction both play an important role and most methods aren’t able to predict non-Watson-Crick pairs in the RNAs. Overall, DeepFoldRNA has the best prediction followed by DRFold as the second best method. Finally, we also suggest possible mitigations to improve the quality of the prediction for future method development.

## Introduction

RNA molecules are essential players in various cellular processes, extending beyond their initial role as passive carriers of genetic information . Their diverse functions, including gene expression regulation, enzymatic activity, and regulatory mechanisms, have highlighted the importance of understanding RNA at a structural level [1]. The three-dimensional (3D) structure of RNA plays a critical role in determining its function [2]. Unlike DNA, RNA molecules can act as single-stranded entities, folding into intricate, dynamic architectures that dictate their biological activity and interactions with other molecules [3]. The 3D structure of RNA is vital for comprehending its diverse functions as the specific spatial arrangement of bases, loops, and stems within RNA molecules influences their ability to interact with proteins, small molecules, and other RNAs [4,5]. This structural organization enables RNA to perform critical tasks, such as serving as ribozymes (catalytic RNA molecules) or regulating gene expression by binding to target mRNA sequences [6,7]. In some cases, small changes in RNA’s 3D structure can significantly impact its function, making accurate structural prediction crucial for understanding RNA biology [1,8].

While there are experimental methods, such as X-ray crystallography, Nuclear Magnetic Resonance (NMR) spectroscopy, and cryo-electron microscopy (Cryo-EM), to determine RNA’s 3D structures, these methods are time-consuming, labour-intensive, and expensive [9,10]. As a result, they are often limited to studying only a fraction of the vast number of RNA molecules present in cells [11]. Moreover, RNA structures are inherently more challenging to resolve due to their dynamic nature, where they can adopt multiple conformations under different cellular conditions [12,13]. Given the challenges and limitations of experimental methods, computational approaches have emerged as essential tools in predicting RNA’s 3D structure [14]. These computational methods leverage existing experimental data, principles of physics, and statistical analyses combined with machine learning to generate plausible models of RNA structures rapidly and cost-effectively [15]. By simulating the folding process, these techniques can provide valuable insights into the energetics and conformational landscape of RNA molecules [16].

Various computational methods have been developed for RNA structure prediction [14,17]. They can be either ab initio, which try to predict the structure from scratch or template-based, which predict the structure based on information from already known structures. The ab initio approaches generally rely on coarse-grained approach to simulate the folding using molecular dynamics and look for the conformation with the lowest free energy [16,18–26]. The template-based methods include fragment-based-assembly (FA-based), comparative modeling, homology modeling, and threading based approaches, which exploit the sequence and structural similarities between known RNA structures and the target RNA, to predict its 3D structure [27–35]. Some methods combine the coarse-grained folding and fragment-based (FA) modeling approaches [36–39]. However, despite significant progress, the accuracy of computational methods for RNA structure prediction remains relatively modest when compared to the progress in computational protein structure prediction [40]. The problem of RNA 3D structure prediction shares many similarities with protein structure prediction, which has been extensively studied [41]. However, predicting RNA structures is often considered more challenging due to several factors. Unlike proteins, there is a relative paucity of experimentally determined RNA structures in publicly available databases (there are only approximately 1700 RNA-only structures in the Protein Data Bank), and structures of RNAs from majority of the RNA families haven’t been experimentally determined (only 87 of the 2791 RNA families have one or more solved 3D structure) [42,43]. This lack of available data makes it harder to establish reliable templates for comparative modelling [15]. Additionally, RNA structures are more dynamic, and their folding pathways are influenced by a myriad of factors with complicated secondary structures, further complicating the prediction process [44]. In recent years, deep-learning-based methods have shown promise in various bioinformatics tasks, including protein structure prediction [45]. Alphafold has revolutionized the field of computational structural modelling, following which, researchers have started exploring the application of similar deep-learning (DL) methods to tackle the challenges of RNA 3D structure prediction [46–48]. The emergence of the end-to-end sequence-based DL methods have shown promise by directly predicting RNA structures from sequence data. They have the advantage of bypassing the need for explicit template structures or extensive feature engineering. DL models, particularly those based on transformer networks and attention mechanisms, can capture long-range complex dependencies within RNA sequences, which are crucial for accurate structure prediction [43,49–53] .

However, it is worth noting that in the last concluded CASP 15 (Critical Assessment of Techniques for Protein Structure Prediction), AI-based methods did not perform as well in RNA structure prediction task compared to their performance in protein structure prediction tasks [54,55]. This highlights the unique challenges posed by RNA structures. Several factors, such as the diversity of RNA families, the varying lengths of RNA molecules, the specific type of RNA (e.g., tRNA, rRNA, mRNA), the degree of sequence homology, and the quality of multiple sequence alignments (MSA), could contribute to the variance in performance. In this study, we systematically benchmark the latest deep-learning-based methods for RNA structure prediction across multiple datasets. Our goal is to investigate the factors contributing to the performance variance, including RNA family diversity, sequence length, RNA type, and MSA quality.

At the conclusion of our study, we provide valuable insights into how DL-based methods for RNA structure prediction can be improved. The results from our independent benchmarking offer guidance on the selection of the most suitable method for different RNA structure prediction use cases. To provide a comprehensive perspective, we have also included two baseline comparisons to non-DL methods, offering a holistic view of the current landscape of 3D RNA structure prediction. Through this systematic benchmarking, we aim to enhance our understanding of the capabilities and limitations of AI-driven approaches in deciphering the 3D structures of RNA molecules, ultimately contributing to advancements in RNA biology and related fields.

## Methods

### RNA Structure prediction methods

We compiled a list of DL-based methods for RNA structure prediction in the recent literature and implemented them locally on our systems to allow a large-scale comparison for multiple targets (Table 1). We also included two fragment-assembly based methods (non-ML) to allow an overall comparison of deep-learning methods against the traditional methods. The details of the benchmarked methods are provided in the Table 1. We have described the benchmarked methods in details in the supplementary (S1).

**Table 1.** The five deep-learning (DL-based) methods and two fragment-assembly (FA-based) methods included in the benchmarking. The MSA columns indicates whether the DL-based methods include MSA as input and the Secondary structure column indicates whether the method uses secondary structure as input. The methods used to predict the MSA and secondary structure are provided in the parenthesis in each column respectively.

| Method | Type | MSA | Secondary structure | Year |
| --- | --- | --- | --- | --- |
| **RosettaFold2NA** | Deep learning (ML) | Yes (rMSA lite) | No | 2022 |
| **DeepFoldRNA** | Deep learning (ML) | Yes (rMSA) | Yes (PETfold) | 2022 |
| **DRFold** | Deep learning (ML) | No | Yes (PETfold + RNAfold) | 2022 |
| **RhoFold** | Deep learning (ML) | Yes (blastn) | No | 2022 |
| **trRosettaRNA** | Deep learning (ML) | Yes (rMSA + Infernal) | Yes (SPOT-RNA) | 2022 |
| **RNAComposer** | Fragment-assembly (non-ML) | - | Yes (RNAfold) | 2012 |
| **3DRNA** | Fragment-assembly (non-ML) | - | Yes (RNAfold) | 2012 |

### Datasets

#### CASP15 RNA

In the 2022 edition of CASP, RNA structure prediction was introduced as a prediction category for the first time [56]. RNA-puzzles experiment has been a long-running CASP-style experiment for RNAs from 2010 to 2021 during which the organizers evaluated 22 RNA-Puzzles challenge [57]. In 2022, RNA-puzzles joined forces with CASP to expand the target and predictors base and stimulate interest in RNA structure prediction field within the protein prediction community. The integration of RNA structure prediction into CASP reflects the growing recognition of the importance of RNA in cellular processes and the need for effective computational tools to predict RNA structures reliably. There were 12 targets selected this year which covered natural RNAs, synthetic RNAs, and RNA-protein complexes (Table 2). We selected this dataset as most of these RNAs have not been seen by the methods that we are benchmarking. Generally, it’s much more difficult to predict the structure of RNA-protein complexes and synthetic RNAs because most of the prediction methods are only trained on single chain RNAs and synthetic RNAs don’t have homologous sequences in the RFam database [58] so the MSA isn’t very informative.

**Table 2.** The targets included in the CASP15 dataset. There are 13 targets in total, out of which four are synthetic and rest are natural RNAs. The difficulty of the targets is taken as estimated by the CASP15 organizers. Four of the targets are X-ray crystallography structures while the remaining nine are Cryo-EM structures. R1189 and R1190 are protein-RNA complex structures which are generally much harder to predict compared to unbound RNAs.

| Name | Type | Length | Experimental Method (resol Å) | RNA Type | Difficulty |
| --- | --- | --- | --- | --- | --- |
| R1107 | RNA | 69 | X<sup>a</sup> (2.8) | CPEB3 ribozyme | Natural/medium |
| R1108 | RNA | 69 | X (2.2) | CPEB3 ribozyme | Natural/medium |
| R1116 | RNA | 157 | X (3.0) | Cloverleaf RNA | Natural/difficult |
| R1117 | RNA-ligand | 30 | X (2.3) | PreQ1 class I type III riboswitch | Natural/easy |
| R1126 | RNA-ligand | 363 | E<sup>b</sup> (5.6) | Traptamer | Synthetic |
| R1128 | RNA | 238 | E (5.3) | Paranemic Crossover Triangle | Synthetic |
| R1136 | RNA-ligand | 374 | E (4.5) | Apta-FRET | Synthetic |
| R1138 | RNA | 720 | E (5.2) | 6HBC | Synthetic |
| R1149 | RNA | 124 | E (4.7) | SARS-CoV-2 SL5 | Natural/difficult |
| R1156 | RNA | 135 | E (5.8) | BtCoV-HKU5 SL5 | Natural/difficult |
| R1189 | RNA-protein | 173 | E (3.8) | RsmZ-RsmA | Natural/difficult |
| R1190 | RNA-protein | 173 | E (4.6) | RsmZ-RsmA | Natural/difficult |

#### New Dataset

We compiled a new dataset of recently published RNA structures in the PDB database [59] to create a more rigorous test of the prediction capabilities of the selected methods (Table 3). We only included RNAs that have been published after the publication of these tools (late 2022) to ensure that none of the RNAs have been included in the training dataset of these methods. This dataset includes RNAs of different types including structures from single RNAs, RNA-ligand complexes, RNA-protein complexes, and synthetic RNAs thus covering the diversity of the RNA structure prediction field. As the methods we benchmarked can only make prediction on single chain RNAs (except RosettaFold2NA), we only made structure predictions on single RNA sequences while discarding the other chains. Our final dataset had 24 targets out of which 16 were natural RNAs and 8 were synthetic. The length of the target RNAs varies from very small RNAs (only 14 nucleotides long) to much longer RNAs (426 nucleotides long). Most of these recently published RNAs have been determined using Cryo-electron microscopy (14/24) [60], with the remaining ones using X-Ray crystallography (9/24) [61] and one using solution NMR [62]. This set of RNAs should serve as the most updated and stringent benchmarking dataset for existing and newly developed RNA structure prediction methods.

**Table 3:**
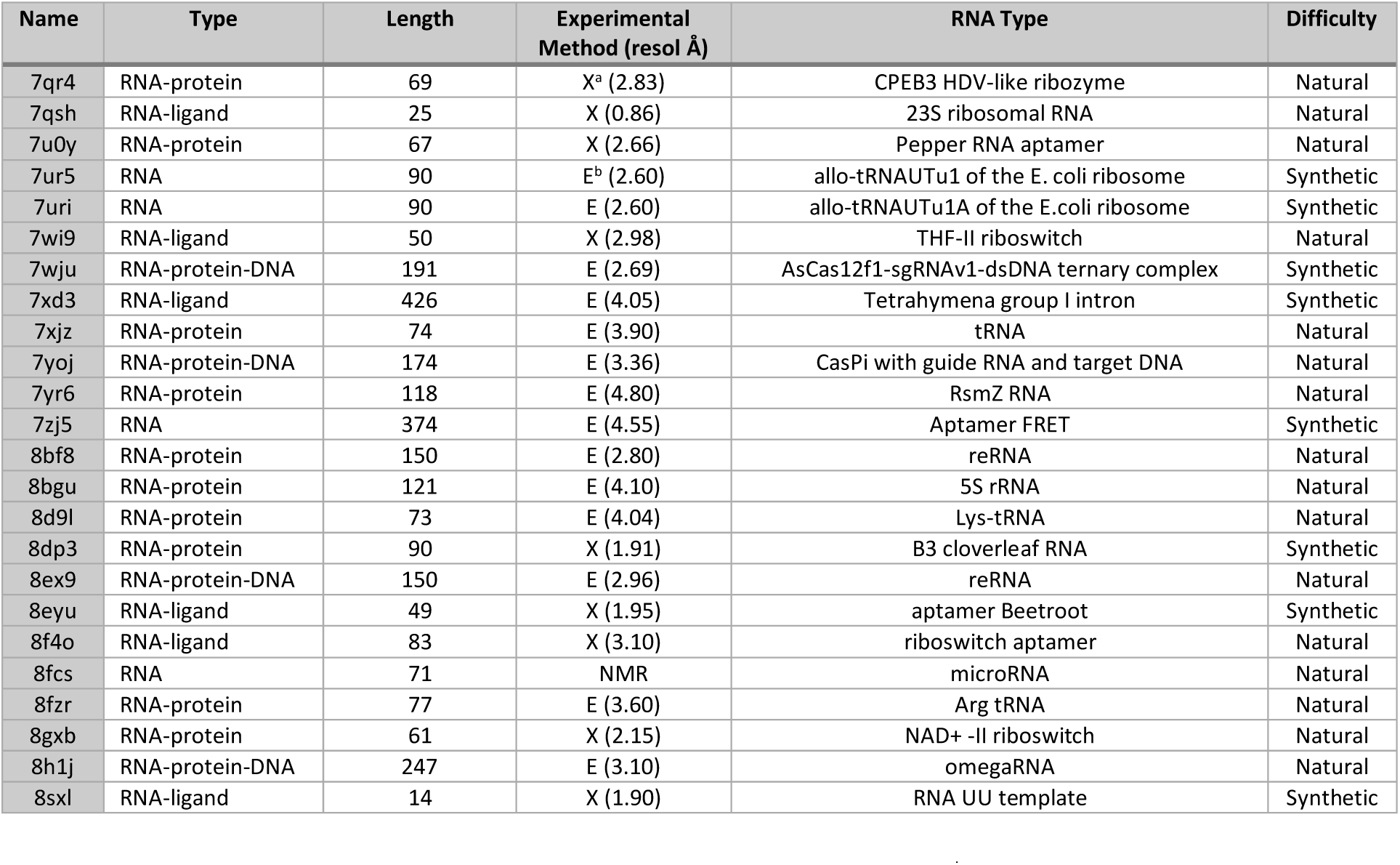
Targets compiled in the new RNA dataset. ^a^X = X-Ray crystallography, ^b^E = Cryo-electron microscopy. All the targets were selected based on their deposition date in the PDB database to make sure that none of the benchmarking-methods contain these targets in their training set.This dataset is well-balanced with 14/24 coming from Cryo-EM and 9/24 coming from X-ray crystallography. 16 of the targets are natural RNAs while 8 are synthetic RNAs.

#### RNA-puzzles dataset

The RNA-Puzzles dataset (Table 4) is a collection of experimentally determined RNA 3D structures that are used in the critical assessment of RNA 3D structure prediction methods, known as the “RNA-Puzzles” experiment [57,63]. RNA-Puzzles is an ongoing collaborative experiment and community effort in the field of structural biology which aims to assess the state of the art in RNA tertiary structure prediction. It is inspired from the CASP experiment for protein structure prediction. The dataset consists of RNA structures whose 3D coordinates have been determined through experimental techniques such as X-ray crystallography, NMR spectroscopy, or cryo-electron microscopy. These experimental structures serve as ground truth for assessing the quality of predicted structures. RNA Puzzles includes RNA structures from various categories, including natural RNA, synthetic RNA, and RNA-protein complexes, where each category presents unique challenges for structure prediction. Since most of the rounds of RNA puzzles were held before 2020 (up to puzzle 24), many of the targets in this dataset might have been part of the training set of the deep-learning methods that we are trying to benchmark, therefore, the benchmarking performance on this dataset could be overestimated. Also, out of the 36 puzzles in our dataset, 35 have been determined using X-ray crystallography, which means that the comparisons with the native structure are more informative about the capabilities of the prediction methods (X-ray crystal structures are more reliable than Cryo-EM or NMR structures). The latest RNA-puzzles targets (published after 2022) are PZ34, PZ37 and PZ38 (PZ39 has already been included in our newly compiled dataset) and these targets should be comparatively more challenging for the prediction methods.

**Table 4:** Targets from the RNA-puzzles dataset. ^a^X = X-Ray crystallography, ^b^E = Cryo-electron microscopy. Most of thes targets were published before the publications of the ML-base methods so it’s highly possible that these are part of the training data of those tools. This will make this dataset the easiest dataset for prediction and the reported performance of most methods might be overinflated on this dataset. Also, 35 out of the 36 targets are X-ray crystallography structures which are known to be more accurate and have better resolution, so the comparison of the predicted model with these native crystallographic structures is more accurate. The puzzle pz34, pz37 and pz38 have been published recently so these targets should be comparatively more challenging for our benchmarking methods.

| Name | PDB Id | Type | Length | Experimental Method (resol Å) | RNA Type | Difficulty |
| --- | --- | --- | --- | --- | --- | --- |
| pz1 | 3MEI | RNA-ligand | 23 | X<sup>a</sup> (1.97) | thymidylate synthase mRNA | Natural |
| pz2 | 3P59 | RNA-ligand | 15 | X (2.18) | RNA Nanosquare | Natural |
| pz3 | 3OWZ | RNA-ligand | 84 | X (2.95) | Glycine riboswitch | Natural |
| pz4 | 3V7E | RNA-Protein | 126 | X (2.80) | SAM-I riboswitch aptamer | Natural |
| pz5 | 4P9R | RNA-ligand | 188 | X (2.70) | Group I intron | Natural |
| pz6 | 4GXY | RNA-ligand | 158 | X (3.05) | Adenosylcobalamin riboswitch | Natural |
| pz7 | 4R4V | RNA-ligand | 185 | X (3.07) | Varkud satellite ribozyme | Natural |
| pz8 | 4L81 | RNA-ligand | 96 | X (2.95) | SAM riboswitch | Natural |
| pz9 | 5KPY | RNA-ligand | 71 | X (2.00) | 5-hydroxytryptophan aptamer | Synthetic |
| pz10 | 4LCK | RNA-Protein | 96 | X (3.20) | T-box riboswitch - tRNA complex | Natural |
| pz11 | 5LYS | RNA-ligand | 57 | X (2.32) | 7SK 5'-hairpin - Gold derivative | Natural |
| pz12 | 4QLM | RNA-ligand | 117 | X (2.72) | ydaO riboswitch | Natural |
| pz13 | 4XW7 | RNA-ligand | 60 | X (2.50) | ZMP Riboswitch | Natural |
| pz14 | 5DDO | RNA-Protein | 58 | X (3.10) | L-glutamine riboswitch | Natural |
| pz15 | 5DI4 | RNA-ligand | 48 | X (2.95) | Hammerhead Ribozyme | Synthetic |
| pz16 | 6Y0Y | RNA-ligand | 27 | X (0.95) | Sarcin Ricin Loop | Synthetic |
| pz17 | 5K7C | RNA-Protein | 58 | X (2.73) | Pistol ribozyme | Synthetic |
| pz18 | 5TPY | RNA-ligand | 71 | X (2.81) | resistant RNA from Zika virus | Natural |
| pz19 | 5T5A | RNA-DNA | 40 | X (2.00) | One Twister Sister ribozyme | Natural |
| pz20 | 5Y85 | RNA-DNA | 71 | X (2.00) | 4-way junctional Twister-Sister Ribozyme | Natural |
| pz21 | 5NWQ | RNA-ligand | 41 | X (1.91) | Guanidine III Riboswitch | Natural |
| pz22 | 6JQ5 | RNA | 82 | X (2.06) | Hatchet Ribozyme | Synthetic |
| pz23 | 6E8U | RNA-ligand | 37 | X (1.55) | Mango-III aptamer | Synthetic |
| pz24 | 6OL3 | RNA-ligand | 112 | X (2.74) | Adenovirus-associated RNA | Natural |
| pz25 | 6P2H | RNA-ligand | 69 | X (2.80) | 2'-dG-II riboswitch | Synthetic |
| pz26 | 6PMO | RNA-ligand | 66 | X (2.66) | T-box riboswitch - tRNA complex | Natural |
| pz27 | 6POM | RNA | 170 | E<sup>b</sup> (4.90) | T-box riboswitch - tRNA complex | Natural |
| pz28 | 6UFM | RNA-Protein | 98 | X (2.82) | T-box riboswitch - tRNA complex | Natural |
| pz29 | 6TB7 | RNA-ligand | 52 | X (2.53) | NAD<sup>+</sup> riboswitch | Natural |
| pz30 | 7BG9 | RNA-Protein | 88 | E (3.80) | Human telomerase RNA | Synthetic |
| pz31 | 7MLX | RNA-Protein | 65 | X (2.09) | PFSE in Sars-CoV-2 | Natural |
| pz32 | 7EOJ | RNA-ligand | 49 | X (1.77) | Pepper Aptamer | Synthetic |
| pz33 | 7ELP | RNA-ligand | 46 | X (2.79) | Xanthine riboswitch | Natural |
| pz34 | 7V9E | RNA-ligand | 68 | X (2.30) | Methyl transferase ribozyme | Natural |
| pz37 | 8GXC | RNA-Protein | 61 | X (2.50) | NAD<sup>+</sup> II riboswitch | Natural |
| pz38 | 8HB8 | RNA-Protein | 55 | X (2.30) | NAD<sup>+</sup> II riboswitch | Natural |

### Assessment Metrics

We used root-mean-square deviation (RMSD) of atomic positions as the metric for quantifying similarity between the modeled and the native structure. RMSD basically measures the average distance between the corresponding atoms of the equivalent residues of two superimposed structures [64]. The predicted 3D models might have different number of residues/atoms than the native structure, therefore we used the RNA-puzzle toolkit [65] to normalize the predicted structures to the native structures. Sometimes the native PDF file might have some part of the structure or residues missing, so we only compared the common residues in the model and native structures. Additionally, hydrogen atoms were removed from the structures. The RMSD between two structures v and w, with n atoms is defined as:

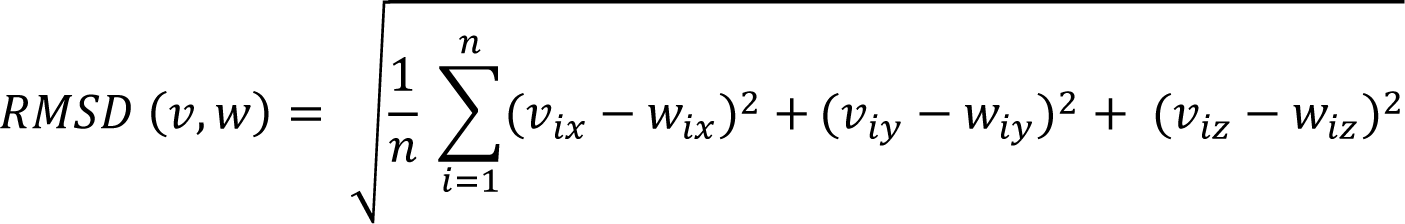

Here, 𝑖𝑥, 𝑖𝑦, 𝑖𝑧 denote the cartesian co-ordinates of the 𝑖^𝑡ℎ^atom. We calculated the all-atom RMSD for comparing the structures i.e. using all the atoms in the residues (except Hydrogens) instead of just heavy atoms or the backbone atoms. While root mean square deviation (RMSD) measurements are commonly employed to assess structural similarity, they may not provide a comprehensive evaluation of the accuracy of predicted RNA 3D structures in comparison to the native structure. For example, two RNA structures may exhibit similar all-atom RMSD values but differ in their base orientations, making RMSD alone an ambiguous criterion. To address this limitation, various metrics and assessment tools have been introduced [66]. These additional measures aim to provide a more nuanced and informative assessment of RNA structural predictions. Interaction network fidelity (INF) is a metric that compares the Watson-Crick (WC) and non-Watson-Crick (nWC) base pairs in the models to the native structure.

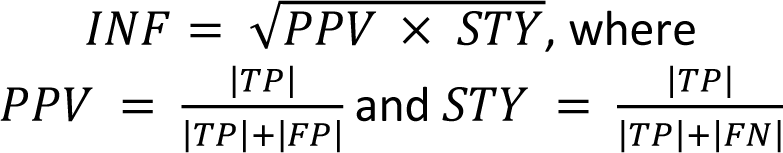

Here TP, FP, and FN are defined based on the absence of base-pairs in the native and modelled structures. We also calculated the Template Modeling score (TMscore) of the predicted models, which is a well-known metric in computational structural biology for measuring the quality of predicted models and is independent of the size of the target unlike RMSD [67].

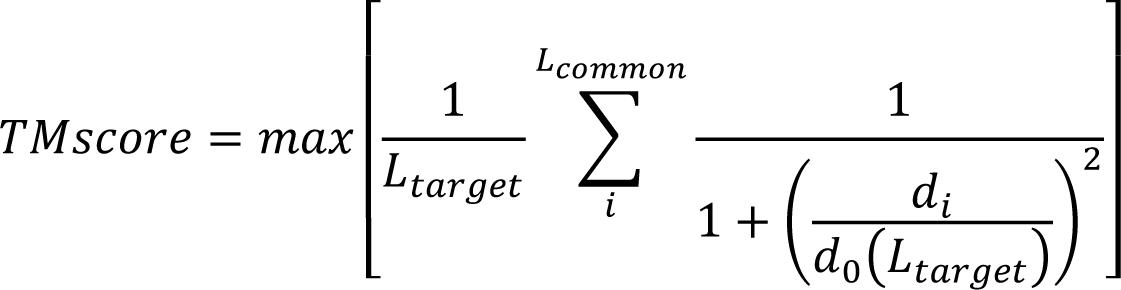

Here, 𝐿_𝑡𝑎𝑟𝑔𝑒𝑡_ is the length of the target(native) RNA, and 𝐿_𝑐𝑜𝑚𝑚𝑜𝑛_ is the number of residues common in both the template(model) and target(native) structures. 𝑑_𝑖_ is the distance between 𝑖^𝑡ℎ^pair of residues in the model and native structures, and 𝑑_0_(𝐿_𝑡𝑎𝑟𝑔𝑒𝑡_) = 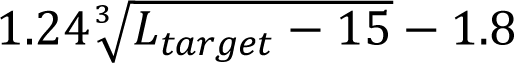 is a scaling factor which approximates the average distance of corresponding residue pairs of random related proteins (making TMscore independent of protein length). TMscore provides a score between 0 and 1, with 1 being an exact match to the native and 0 being the lowest. Generally, a TMscore <0.17 means that the structures are indistinguishable from random pairs, while a score >0.45 suggests that the structures have the same global topological fold. As TMscore gives larger weights to smaller distance errors, it makes the score more sensitive to the global fold similarity unlike RMSD, which is more sensitive to local structural variations. We included an additional metric to measure the fraction of native contacts recovered in the modelled structures [68]. This Native Contact Fraction metric calculates the fraction of residue contacts (a contact is defined as two non-contiguous residues occurring within a distance cut-off of 5 Å) that are also present in the modelled structures.

## Results

### Results on the CASP15 dataset

The native structures for all the 13 targets in this dataset were not available to us, so we only compared our models for seven targets. The average RMSD of the DRFold predicted models was the lowest (RMSD = 27.38 Å), while the 3DRNA models had the highest RMSD 34.76 Å overall (Table 5; Fig 1a). Notably, the best models from the CASP15 methods were much better than the ones in our benchmarking (Table 5). The RMSDs for natural targets (R1107 and R1108) were much lower than that for the other targets for all the methods, and the RMSDs for both the synthetic and RNA-protein complexes were much higher (Fig 1a). Same was true for other metrics as well (Fig 1b; Fig S1). We selected two targets (R1107 and R1136) as examples for visualizing good and poor quality predictions by showing the 3D structural alignment of their models with the native structure. The models predicted by most methods for R1107 are predicted reasonably well and we see a good alignment with the native structure (Fig 2). All the models for R1136 were of poor quality as it is a very long synthetic RNA and therefore the structural alignment wasn’t good at all (Fig 3).

**Table 5.** Comparison of RMSD values of the predicted models by the various methods to native structure for targets in the CASP15 dataset. DRFold has the lowest average RMSD of 27.38 Å, which is slightly better than that of DeepFoldRNA (28.83 Å). ‘best_rmsd_ours’ column denotes the rmsd of the best model predicted by our benchmarking methods and ‘best_rmsd_casp’ column contains the rmsd of the best model from all the methods participating in CASP15. Looking at the ‘best_rmsd_ours’ column clearly shows that we are only able to predict models of reasonably well quality for only two of the seven targets(r1107 and r1108). df: DeepFoldRNA, drfold: DRFold, rf: RosettaFold2NA, rhofold: RhoFold, rnac: RNAComposer, trr: trRosettaRNA, 3drna: 3dRNA

|  | df | drfold | rf | rhofold | rnac | trr | 3drna | best_rmsd_ours | best_rmsd_casp |
| --- | --- | --- | --- | --- | --- | --- | --- | --- | --- |
| **r1107** | 6.19 | 18.3 | 9.58 | 7.79 | 19.12 | 13.33 | 22.36 | 6.19 | 4.5 |
| **r1108** | 5.67 | 9.24 | 11.25 | 6.49 | 19.64 | 5.57 | 23.25 | 5.67 | 4.5 |
| **r1126** | 49.09 | 41.45 | 57.12 | 43.32 | 54.68 | 44.45 | 39.84 | 39.84 | 4.3 |
| **r1128** | 24.71 | 26.39 | 24.6 | 20.67 | 26.98 | 19.52 | 38.05 | 19.52 | 7.2 |
| **r1136** | 37.26 | 50.08 | 53.88 | 55.94 | 42.49 | 38.27 | 43.72 | 38.27 | 7.8 |
| **r1138** | 60.74 | NA | 63.71 | 64.38 | NA | 49.02 | 51.28 | 49.02 | 16.6 |
| **r1189** | 23.51 | 23.01 | 25.25 | 22.68 | 25.46 | 35.22 | 29.72 | 23.01 | 16 |
| **r1190** | 23.5 | 23.22 | 25.38 | 23.13 | 25.48 | 36.85 | 29.89 | 23.13 |  |
| **Average** | **28.83** | **27.38** | **33.85** | **30.55** | **30.55** | **30.28** | **34.76** |  |  |

**Fig 1.**
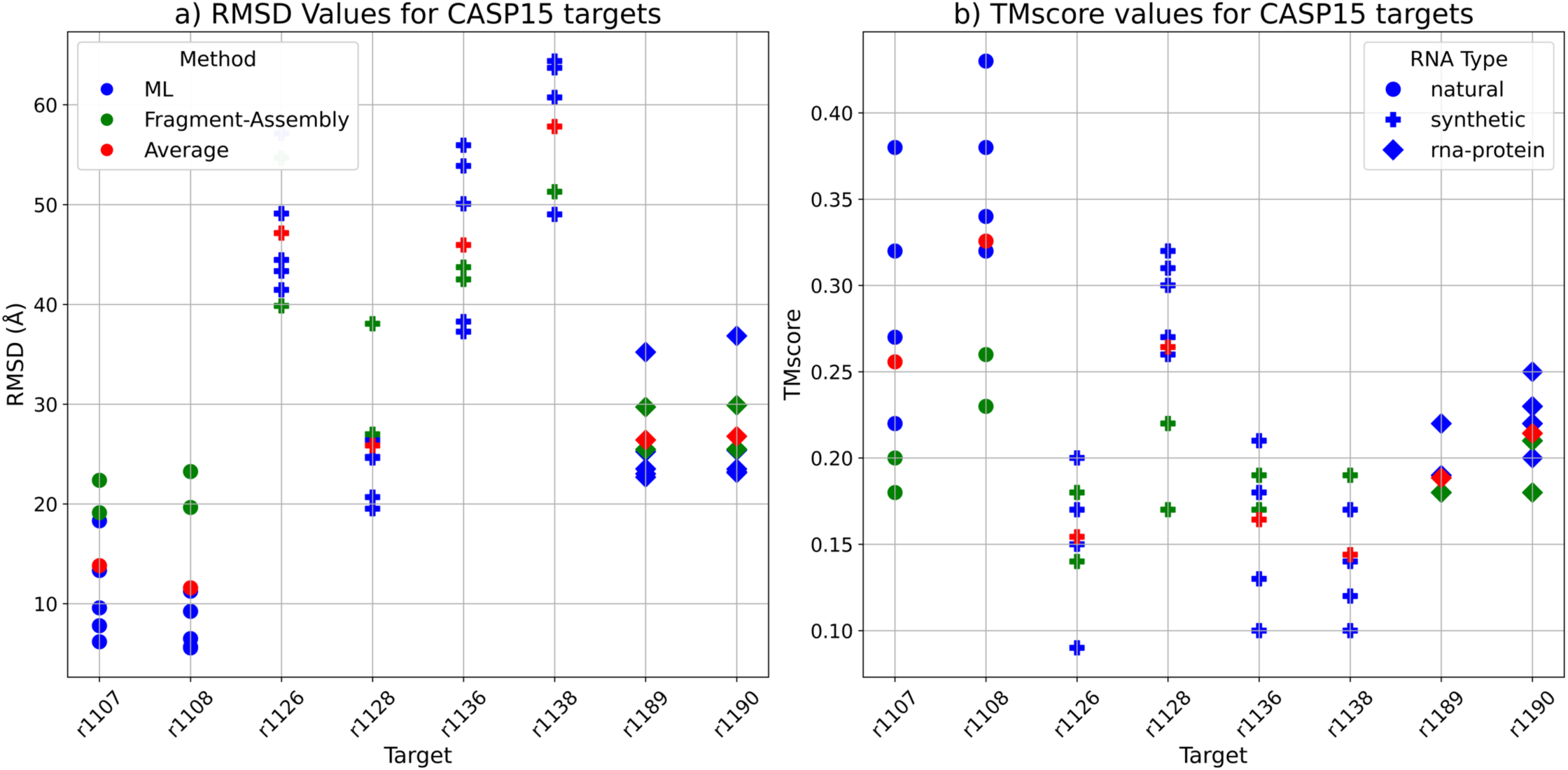
The RMSD and TMscore comparison for the RNA targets in the CASP15 dataset. Models predicted by Machine-Learning-based (ML-based) methods are coloured in blue, the ones predicted by Fragment-Assembly-based (FA-based) methods are in green and the average RMSD of all models for each target is in red. The shape of the points is based on the RNA type with circle denoting a natural RNA, + denoting a synthetic RNA and a square denoting an RNA-protein complex **a)** Plot showing the RMSD values in Å for the seven targets in the CASP15 dataset. For natural RNAs (r1107 and r1108), the ML-based methods (in blue) have much lower RMSD than the average (in red) and the FA-based methods (in green). The best model for each target is the one usually predicted by a ML-method (except for R1126 which is a synthetic target with a length of 363 nucleotides). The average RMSD for most synthetic targets is higher than the natural and RNA-protein complex targets. **b)** Plot showing the TMscores for the predicted models for each target. TMscore for natural targets (r1107 and r1108) is much higher compared to the synthetic and RNA-protein targets. For the natural targets, ML-predicted models have higher TMscore than the average and the FA-predicted models. Model with the best TMscore for each target is one predicted by a ML-based method (except for r1138 which is a very long synthetic RNA of 720 nucleotides)

**Fig 2.**
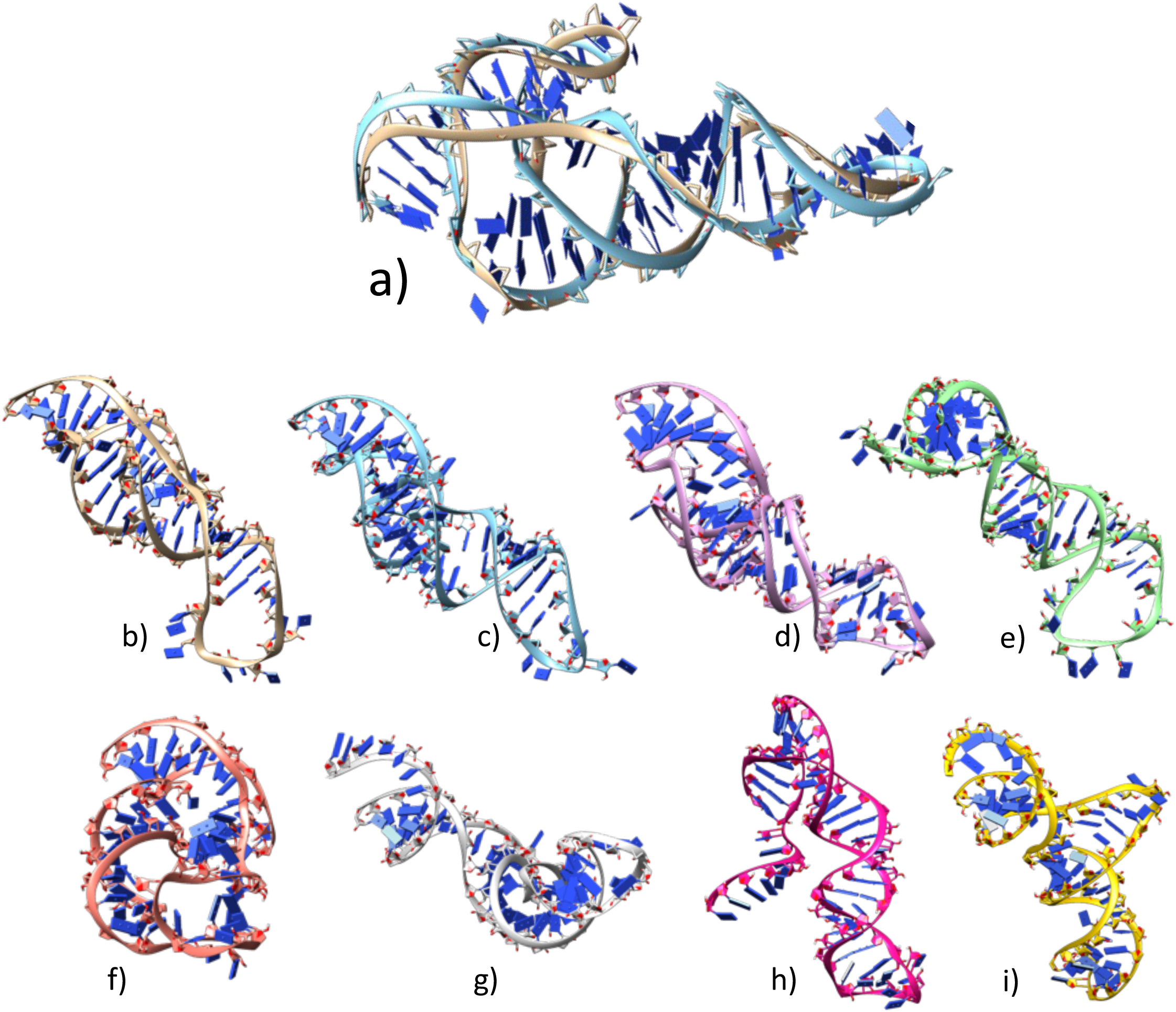
Native and predicted models for CASP target R1107. All the structures were aligned together against the native(b) and then tiled separately to visualize. **a)** Superimposition of native structure (in beige colour) to the best model (DeepFoldRNA, cyan colour); RMSD=6.19 Å **b)** Native structure **c)** DeepFoldRNA model **d)** RhoFold model; RMSD=7.79 Å **e)** RosettaFold2NA model; RMSD=9.58 Å **f)** trRosettaRNA model; RMSD=13.35 Å **g)** DRFold model; RMSD=18.30 Å **h)** RNAComposer model; RMSD=19.12 Å **i)** 3DRNA model; RMSD=22.54 Å

**Fig 3.**
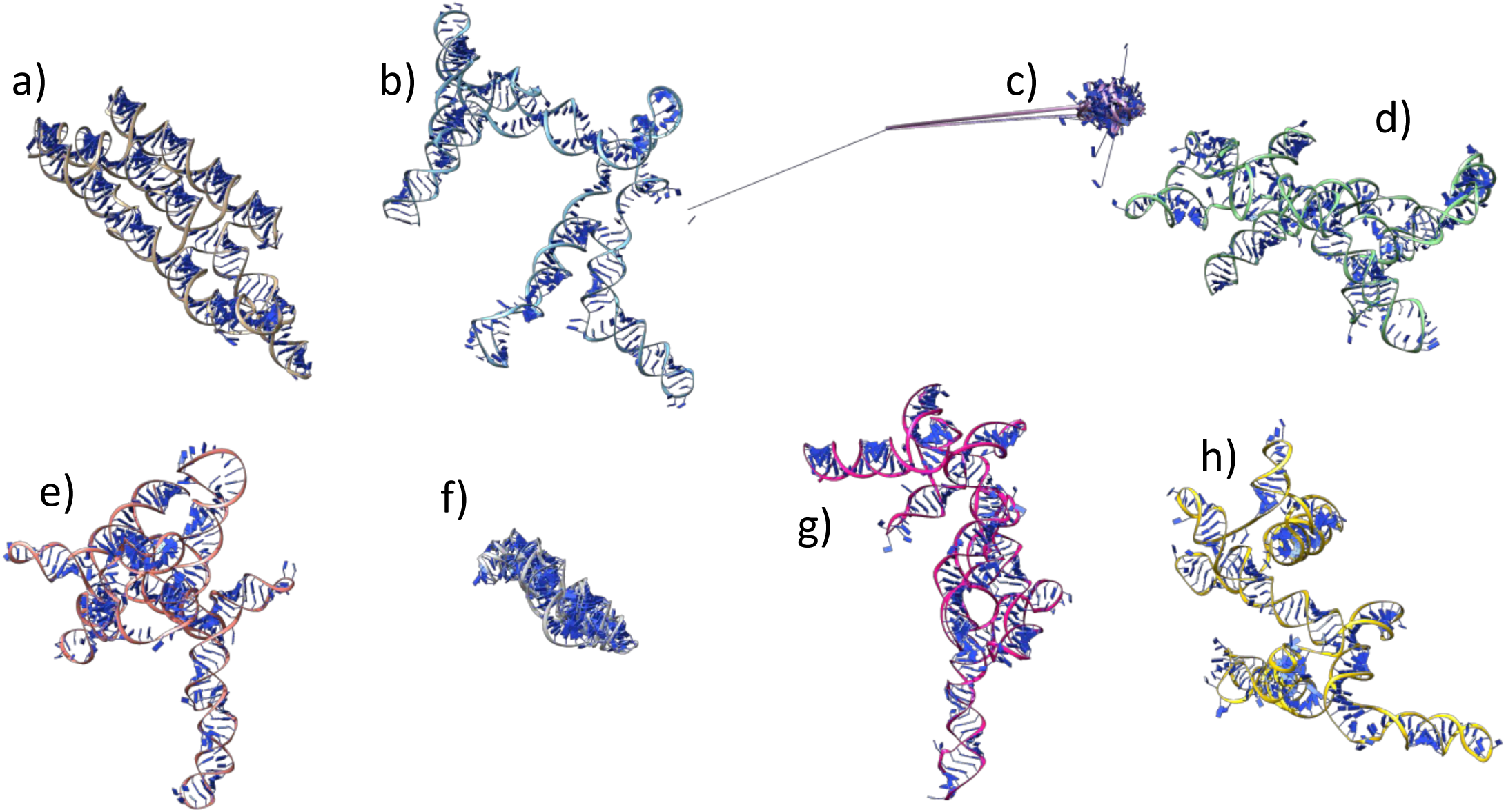
Native and predicted models for CASP target R1136. All the structures were aligned together against the native(a) and then tiled separately to visualize. **a)** Native structure **b)** DeepFoldRNA model; RMSD=37.26 Å **c)** RhoFold model; RMSD=55.94 Å **d)** RosettaFold2NA model; RMSD=53.88 Å **e)** trRosettaRNA model; RMSD=38.27 Å **f)** DRFold model; RMSD=50.08 Å **g)** RNAComposer model; RMSD=42.49 Å **h)** 3dRNA model; RMSD=43.72 Å

### Results on the New dataset

On this dataset, the DeepFoldRNA has the lowest average RMSD overall, with DRFold being the close second (Table 6). The fragment-assembly based methods had the highest RMSD overall with the worst prediction performance (Table 6; Fig 4). We also observed that generally the natural targets had lower RMSD (and higher TMscore) when compared to the synthetic and RNA-protein complex targets (Fig 4). The ML-based methods again had a better performance than the FA-based methods with the best model (based on RMSD and TMscore) for each target being always the one predicted by a ML-based method. This was true for other metrics as well (Fig S3).

**Table 6.**
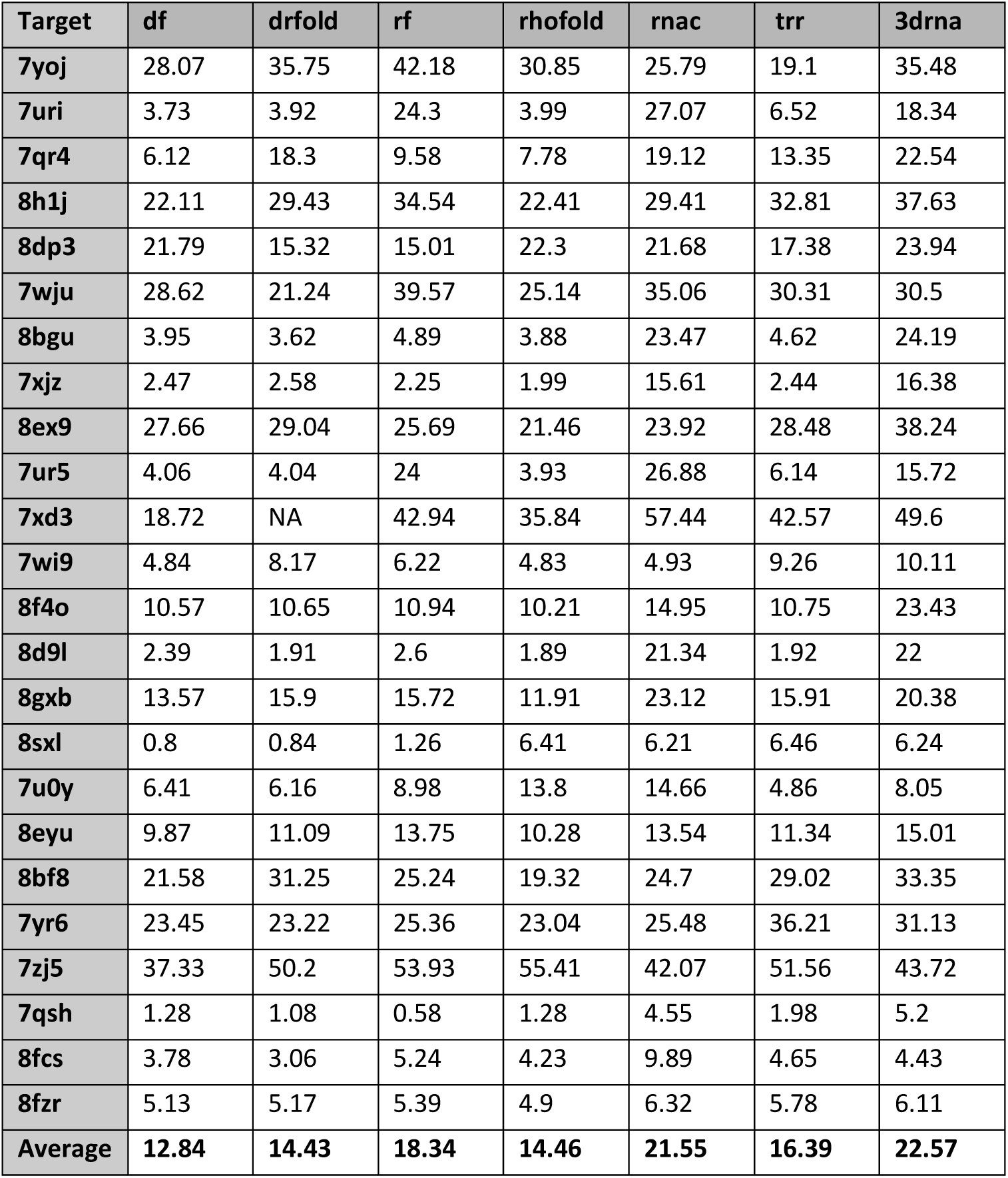
Comparison of RMSD values of the predicted models to the native structure for targets in the newly compiled dataset. DeepFoldRNA(df) has the lowest average RMSD (12.84 Å) on this dataset. RMSD for the fragment-assembly-based methods (3DRNA and RNAComposer) is much higher than that of the deep-learning-based methods. df: DeepFoldRNA, drfold: DRFold, rf: RosettaFold2NA, rhofold: RhoFold, rnac: RNAComposer, trr: trRosettaRNA, 3drna: 3dRNA

**Fig 4.**
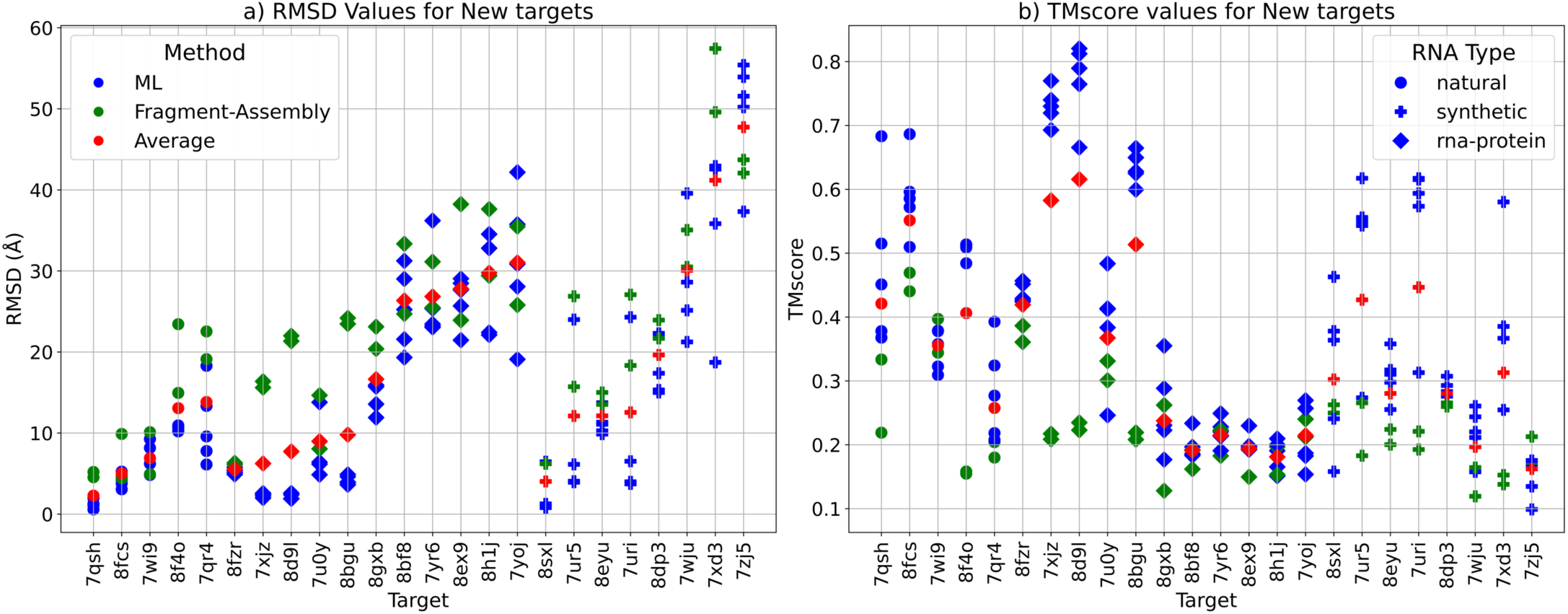
The RMSD and TMscore comparison for the RNA targets in the New dataset. Models predicted by Machine-Learning-based (ML-based) methods are coloured in blue, the ones predicted by Fragment-Assembly-based (FA-based) methods are in green and the average RMSD of all models for each target is in red. The shape of the points is based on the RNA type with circle denoting a natural RNA, + denoting a synthetic RNA and a square denoting an RNA-protein complex **a)** Plot showing the RMSD values in Å for the targets in the New dataset. For most targets, the ML-based methods (in blue) have much lower RMSD than the average (in red) and the FA-based methods (in green). The average RMSD for most synthetic targets is higher than the natural and RNA-protein complex targets. **b)** Plot showing the TMscores for the predicted models for each target. TMscore for almost all targets for ML-methods (in blue) is higher compared to the Average(in red) and FA-based methods (in green). Model with the best TMscore for each target is one predicted by a ML-based method. The average TMscore for most synthetic targets is higher than the natural and RNA-protein complex targets.

### Results on the RNA-Puzzles dataset

The average results on this dataset are much better than the other datasets. DRFold has the best performance (lowest average RMSD) on this dataset followed closely behind by DeepFoldRNA as the second best. The other ML-based methods also have a comparatively lower average RMSDs on this dataset compared to the others. However, the performance of the FA-based methods lags far behind the ML-based methods (Table 7). We also see that on pz34, pz37 and pz38 targets (more challenging as they were definitely not included in the training set of ML-based methods), the performance of our methods was much worse (except for RhoFold which has 2.65 Å model for pz34 because it was published in early 2022 and possibly includes it in its training set). Surprisingly, unlike the other datasets we didn’t see a much better average performance for natural targets compared to synthetic and RNA-complex targets on this dataset (Fig 5). The performance of the methods based on other metrics was also a lot better on this dataset (Fig S4).

**Table 7.**
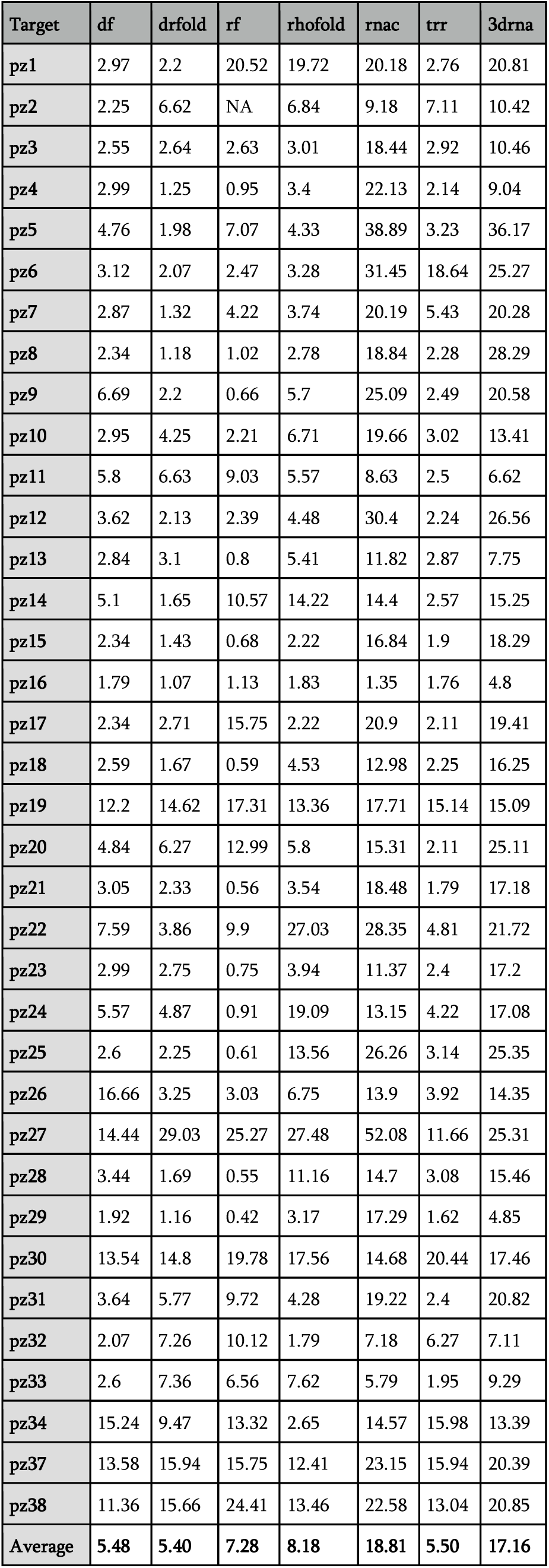
Comparison of RMSD values (in Å) of the predicted models to the native structure for targets in the newly compiled dataset. DRFold has the lowest average RMSD (5.40 Å) on this dataset. RMSD for the fragment-assembly-based methods (3DRNA and RNAComposer) is much higher than that of the deep-learning-based methods. df: DeepFoldRNA, drfold: DRFold, rf: RosettaFold2NA, rhofold: RhoFold, rnac: RNAComposer, trr: trRosettaRNA, 3drna: 3dRNA

**Fig 5.**
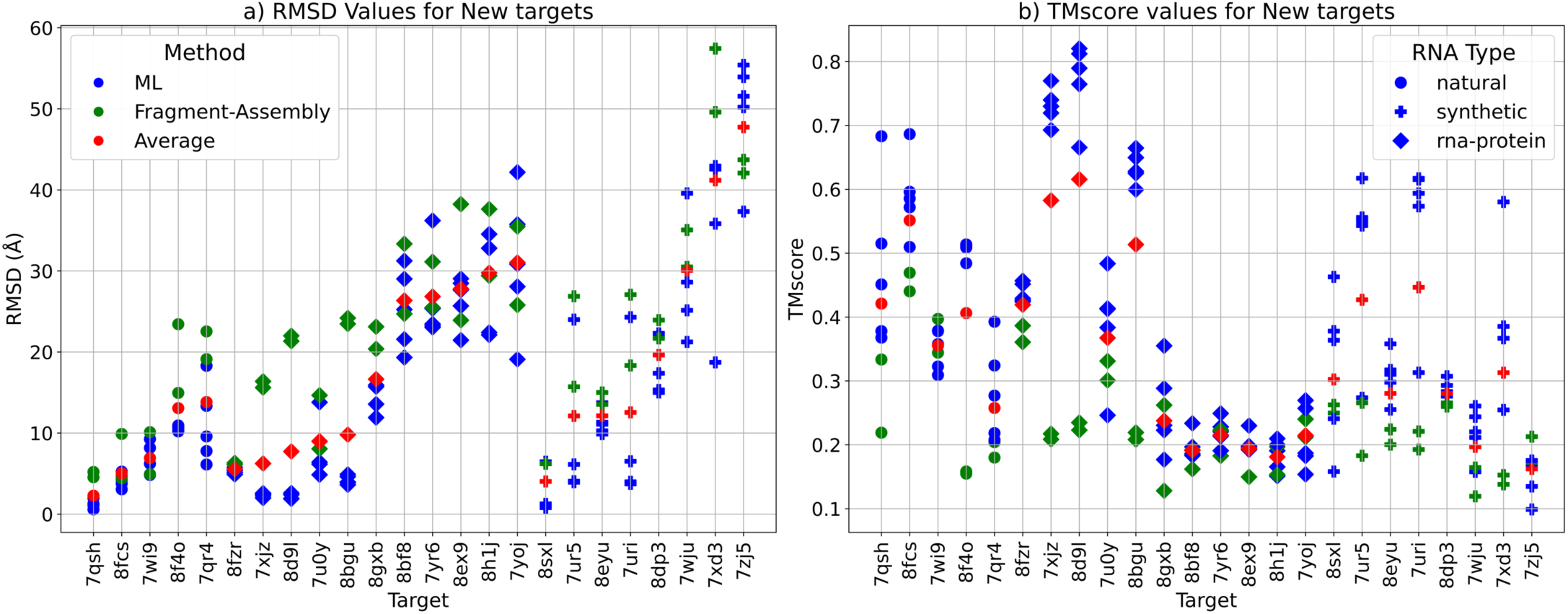
The RMSD and TMscore comparison for the RNA targets in the RNA-puzzles dataset. Models predicted by Machine-Learning-based (ML-based) methods are coloured in blue, the ones predicted by Fragment-Assembly-based (FA-based) methods are in green and the average RMSD of all models for each target is in red. The shape of the points is based on the RNA type with circle denoting a natural RNA, + denoting a synthetic RNA and a square denoting an RNA-protein complex. On average, the performance of most methods on this dataset is much better than on CASP15 or the New dataset, possibly because many targets might have been part of the training set of the ML-methods and also many homologous structures for these targets are available in the PDB. The ML-based methods have the best quality models (low RMSD and High TMscore) and the FA-based methods have the lowest quality models for most targets in this dataset **a)** Plot showing the RMSD values in Å for the targets in the CASP dataset. For most targets, the ML-based methods (in blue) have much lower RMSD than the average (in red) and the FA-based methods (in green). **b)** Plot showing the TMscores for the predicted models for each target. TMscore for almost all targets for ML-methods (in blue) is higher compared to the Average(in red) and FA-based methods (in green). Model with the best TMscore for each target is always the one predicted by a ML-based method.

### Combined results

We finally looked at the performance of all the methods on the combined dataset i.e. CASP15+New+RNA-Puzzles to get an idea about the generalized performance of the methods (Fig 6). The performance was compared using several metrics including RMSD, TMscore, Native contact fraction (Ncf), Interaction network fidelity (INF) for all pairs, INF-wc (INF for Watson-Crick pairs), and INF-nwc (INF for non-Watson-Crick pairs). DeepFoldRNA had the lowest median RMSD overall, while 3DRNA had the highest. We also included average prediction as an additional comparison method, which is basically the average of the metric score of the model predicted by all of the seven methods for each target. We observed that all the five deep-learning methods have a better performance than the average prediction and the two fragment-assembly-based methods have a worse performance than the average prediction based on RMSD (Fig 6a), TMscore (Fig 6b), Ncf (Fig 6c) and INF score (Fig 6d). The INF score is essentially a metric quantifying how accurately is the base-pairing of the nucleotides predicted. The INF-wc and INF-nwc score are the only two metrics where any of the ML-based methods have a worse median than any of the FA-based methods (Fig 6e; 6f). Although most of the methods are able to predict the Watson-Crick pairs quite well (Fig 6e), none of the methods are good in predicting non-cannonical base pairs as the median RMSD of even the best method for INF-nwc score is only 0.47.

**Fig 6.**
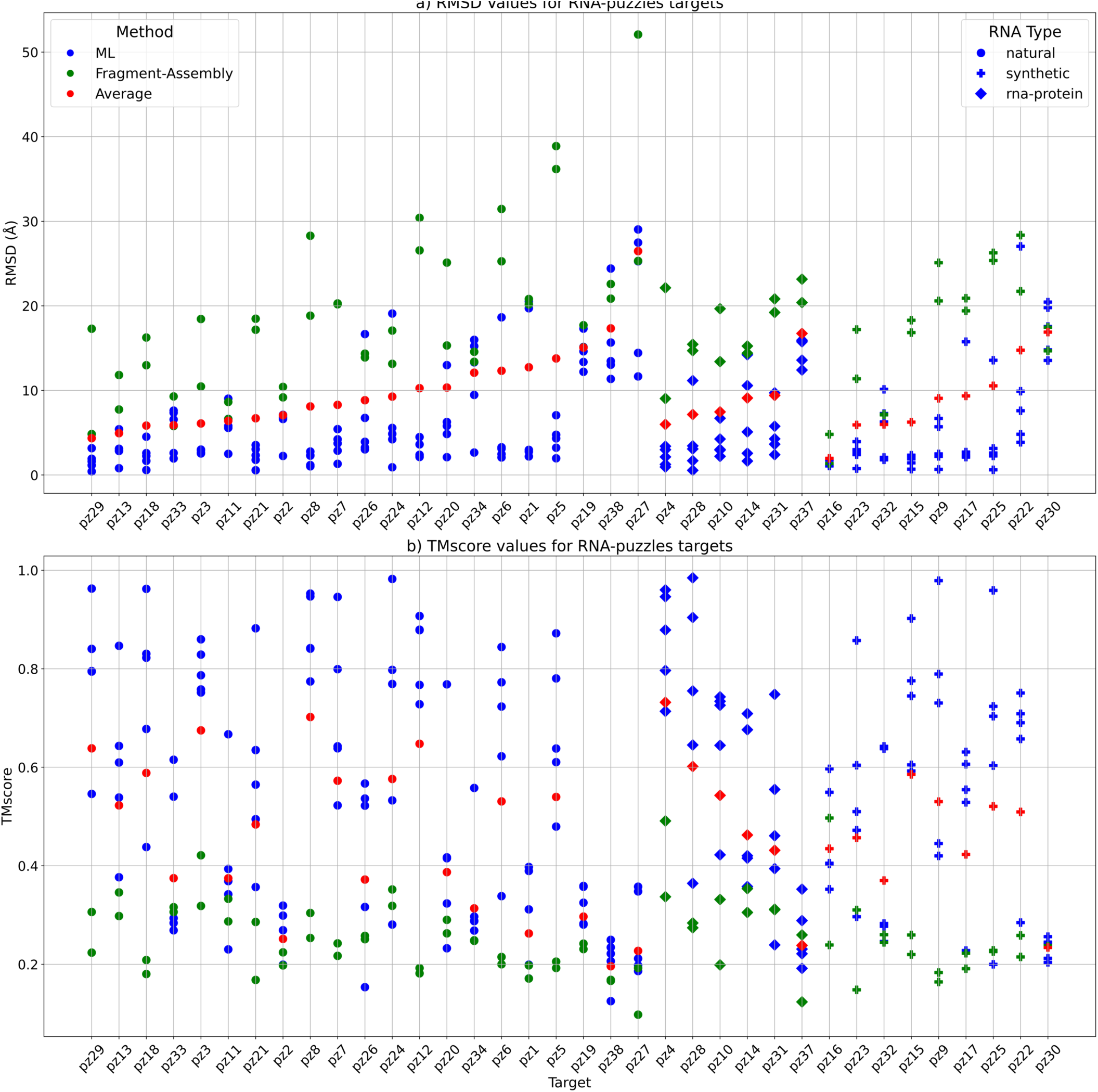
Box and violin plots showing the comparison of all the methods on the combined dataset across multiple metrics. The methods are on the X-axis and the metrics are on the Y-axis. The Average plot (in grey) is the average of the models predicted by all the methods for a particular target. The median values are labelled with blue text and the whiskers denote the interquartile range **a)** RMSD distribution of the predicted models by various methods. DeepFoldRNA has the lowest median RMSD (5.12 Å). **b)** TMscore distribution for the various methods. DeepFoldRNA and DRFold have the highest median TMscore **c)** Native contact fraction (ncf) of the predicted models by the various methods. DRFold has the highest ncf of 0.75. **d)** INF score for the various methods. DeepFoldRNA has the highest median INF score (0.81) **e)** INF-wc score (Watson-Crick pairs) for the various methods. DeepFoldRNA has the highest median score of 0.92. Most methods have a really good INF-wc score (>=0.8 for most methods) indicating that the cannonical Watson-Crick pairs are predicted quite accurately by most methods. Interestingly, for the first time for a metric, a ML-based method i.e. trRosettaRNA has a median score lower than the medians of the Average prediction or FA-based methods (3DRNA, RNAcomposer). This could be because the secondary structure prediction method used by trRosettaRNA might not be as accurate as others. **f)** INF-nwc score (non-Watson-Crick pairs) for the various methods. DeepFoldRNA and RosettaFold2NA have the highest median RMSD of 0.47. None of the methods even have a median score higher than 0.5 indicating that none of the the methods are very good at predicting non-cannonical base pairing. Interestingly, again this time a ML-based method (DRFold) has a lower median score than the Average as well as median score of RNAComposer (an FA-based method). Usually, DRFold has been the second-best method and close to DeepFoldRNA on most metrics, so this discrepancy might be explained because of its non-reliance on MSA as input to predict the structure, as all other ML-based methods use MSA and they are able to predict non-Watson-Crick pairs more accurately.

We can also look at the performance of all the methods by looking at the fraction of correctly predicted targets out of all targets. A predicted model is termed as a correct prediction, if the RMSD/TMscore of the predicted model is less/more than a certain RMSD/TMscore cut-off. Generally in RNA structure prediction field, for RMSD, a model within 5 Å is considered a very good prediction and even 10 Å is considered acceptable, while for TMscore, a score higher than 0.4 to 0.45 indicates a model with similar fold to the native structure. We used RMSD cut-offs of 2.5 Å (almost near-native predictions) to 15 Å with intervals of 2.5 Å and TMscore cut-offs of 0.2 to 1. At 5 Å cut-off, DeepFoldRNA, DRFold and trRosettaRNA had close to 50% correct predictions (32/65), whereas RosettaFold2NA had 37% (24/65) and RhoFold had 40% (26/65). For the non-ML methods, they are only able to only predict less than 5% of the targets (3/65) correctly at a cut-off of 5 Å and it only increases to around 30% (17/65 for 3dRNA and 21/65 for RNAComposer) on increasing the cut-off to 15 Å (Fig 7a). Similar results were seen when we looked at different TMscore cut-offs with DL-based methods being much better (DeepFoldRNA being the best) than the FA-based methods overall (Fig 7b).

**Fig 7.**
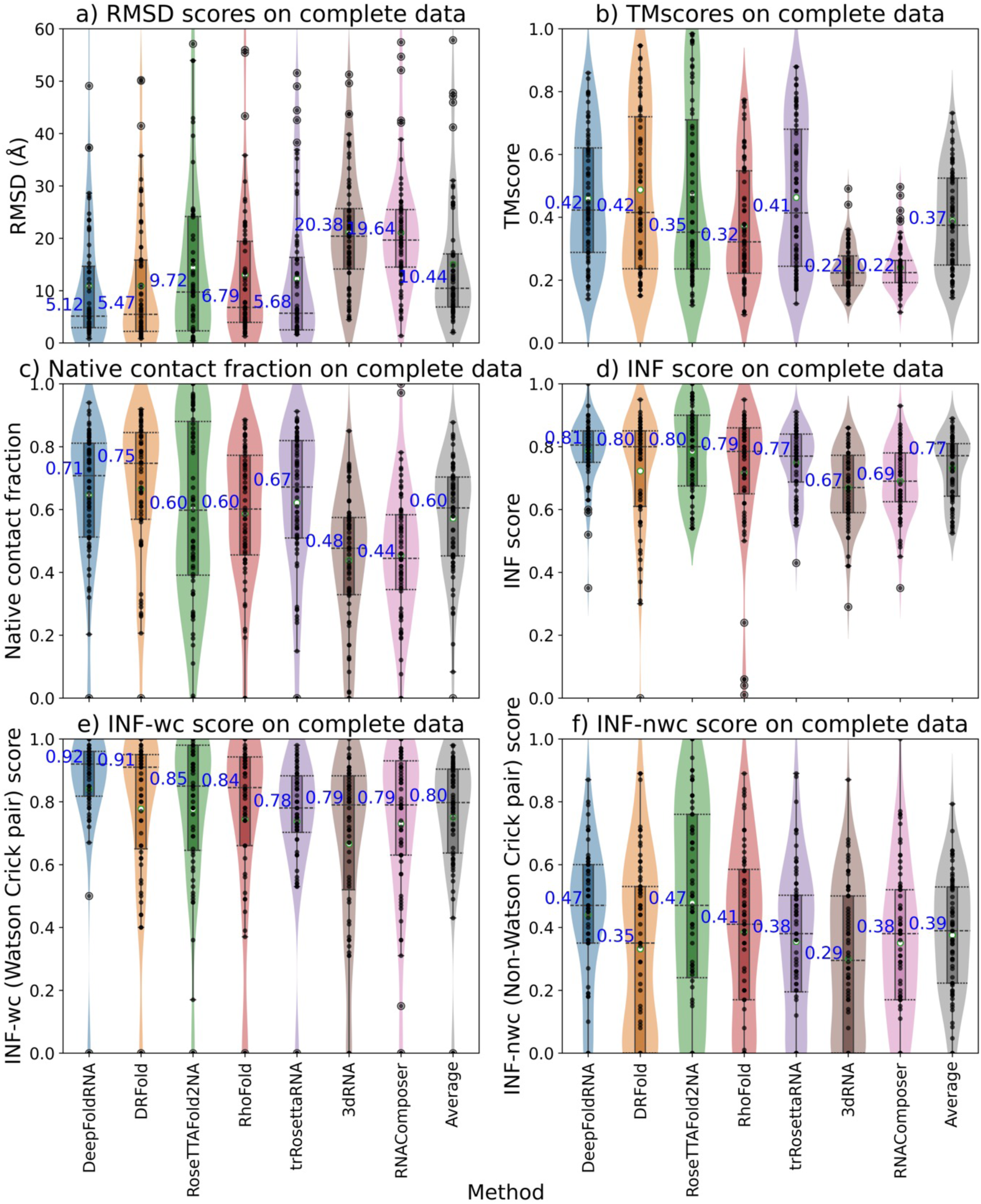
Plots showing the performance of various RNA structure prediction methods at different RMSD and TMscore cut-offs. **a)** At a RMSD cut-off of 5 Å, DeepFoldRNA, DRFold and trRosettaRNA are able to predict 50% of the targets correctly, which increases to 70-75% on increasing the cut-off to 15 Å. However, RNAComposer and 3DRNA are only able to predict 5% of the targets correctly at 5 Å cut-off and even after increasing the RMSD cut-off for correct predictions to 15 Å, they are only able to predict around 30% of the targets correctly. **b)** At a TMscore cut-off of 0.4 most ML-based methods are able to predict ∼50% targets correctly, while FA-based methods are only able to predict <5% targets correctly. On applying a more stringent cut-off of 0.6 the % of correct predictions for the ML-methods drops below 40% while FA-methods aren’t even able to predict a single model with a TMscore higher than 0.6.

### Pairwise comparison of different methods

We created scatter plots of the RMSD of the predicted models for a pairwise comparison of each method against all other methods. In total we created 8×8 scatter plots (5 ML-based, 2 FA-based and one Average prediction). We observed that generally the ML-based methods have a better performance than the Average prediction and the two FA-based methods (Fig 8). Overall the DeepFoldRNA-predicted models have the best accuracy (in RMSD) when compared against models from all other methods. We also created similar scatterplots for all vs all comparison of the methods based on TMscore and there also DeepFoldRNA turned out to be the best method (Fig S5).

**Fig 8.**
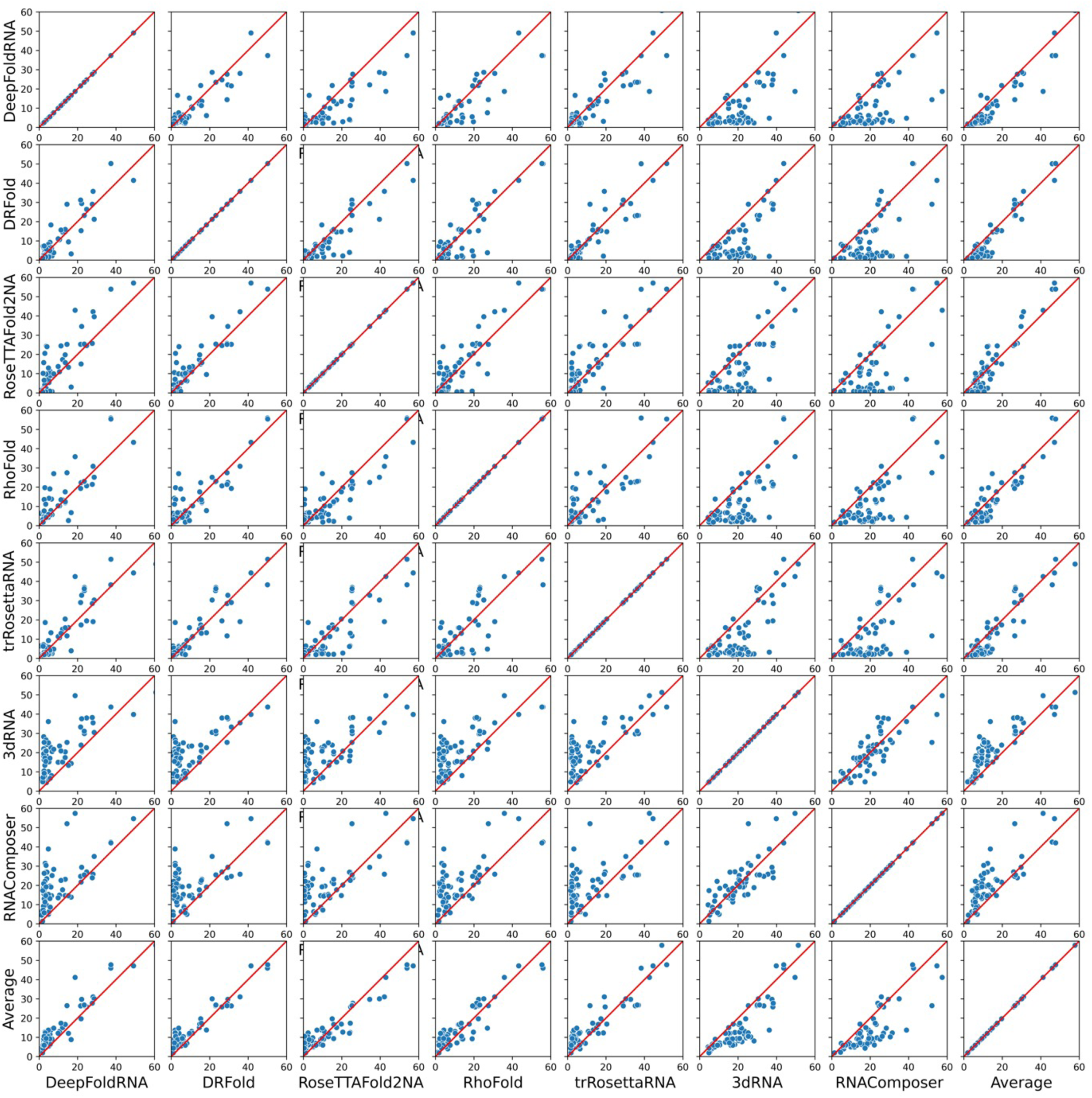
Scatterplot showing the performance comparison of each method against every other method on all the targets. If a point lies on the red-coloured x=y line, it indicates that the RMSD of the predicted model from both the methods is exactly the same i.e. they have similar prediction performance for that target. Points above that line indicate a higher RMSD for the model predicted by the method on the y-axis (i.e. method on the x-axis is better) and points below that line indicate vice-versa. Most of the ML-based methods have a better performance than the average prediction (last row of plots), while the FA-based methods are much worse than the average prediction (Average vs 3dRNA and Average vs RNAComposer plots in the last row). When compared against all other methods using the RMSDs of the predicted models, DeepFoldRNA is the best method followed by DRFold.

### Comparison of ML-based methods with the FA-based methods

We compared the performance of the ML-based methods together against the non-ML or FA-based methods to get an idea about how much better the ML methods are comparatively. The median RMSD of all models predicted by ML-based methods was much lower than the ones predicted by FA-based methods (Fig 9a). We also compared the performance differences based on the datasets (Fig 9b) and RNA type (Fig 9c) and in all cases the ML-based methods were superior to the FA-based methods.

**Fig 9.**
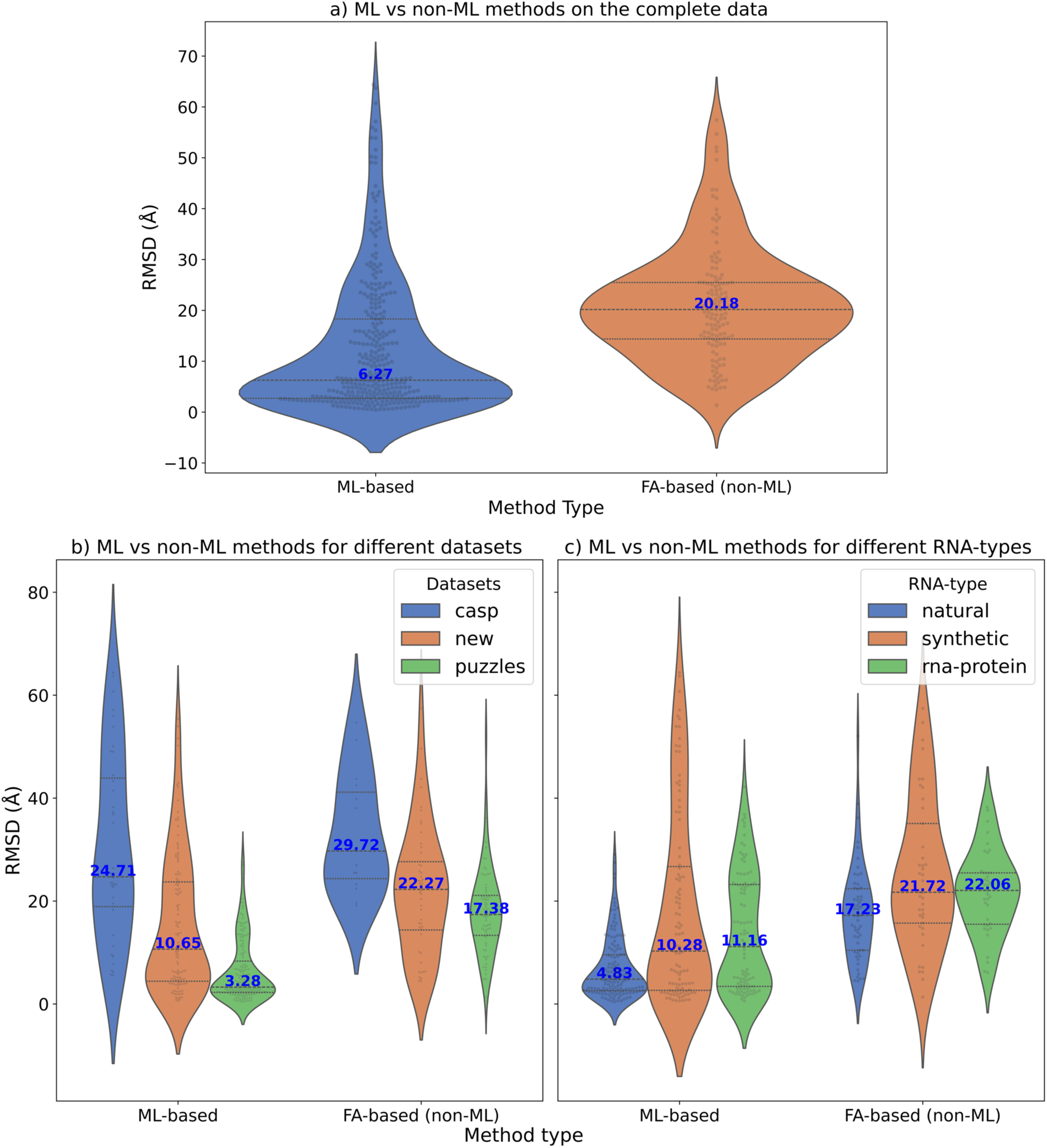
The comparison of ML-based methods to non-ML-based methods. The median values for each violin plot are labelled in blue. The RMSD (in Å) is shown on the y-axis while the x-axis shows the method type (ml or non-ml). **a)** Violin plots showing the distribution of the RMSDs of the predicted models for a comparison between ml (DeepFoldRNA, DRFold, trRosettaRNA, RosettaFold2NA, RhoFold) and non-ml (3dRNA, RNAComposer) methods. The median RMSD of ml methods (6.27 Å) is much lower than non-ml methods (20.18 Å). **b)** Violin plots showing the comparison of ML vs FA-based methods based on different datasets. ML-based methods are clearly better with much lower median RMSD than FA-based methods on the New (10.65 Å vs 22.27 Å) and RNA-puzzles dataset (3.28 Å vs 17.38 Å). ML-based methods are also better than FA-based ones on the CASP15 dataset albeit the difference in median RMSD is not as pronounced (24.71 Å vs 29.72 Å) . **c)** Violin plots showing the comparison of ML vs FA-based methods based on different RNA types. ML-based methods are better with much lower median RMSDs for all RNA types. 4.83 Å vs 17.23 Å for natural, 10.28 Å vs 21.72 Å for synthetic and 11.16 Å vs 22.06 Å for RNA-protein complex targets.

### Effect of different datasets on the prediction accuracy of the methods

The performance of the methods varies depending on the dataset they are benchmarked on. We compared the accuracy of the models (based on RMSD) for all models depending on the dataset and CASP15 was the hardest, New dataset was in the middle and RNA-puzzles was the easiest (fig 10a). We also looked at the performance of each method separately based on the dataset being benchmarked on and we saw similar results (fig 10b; S6).

**Fig 10.**
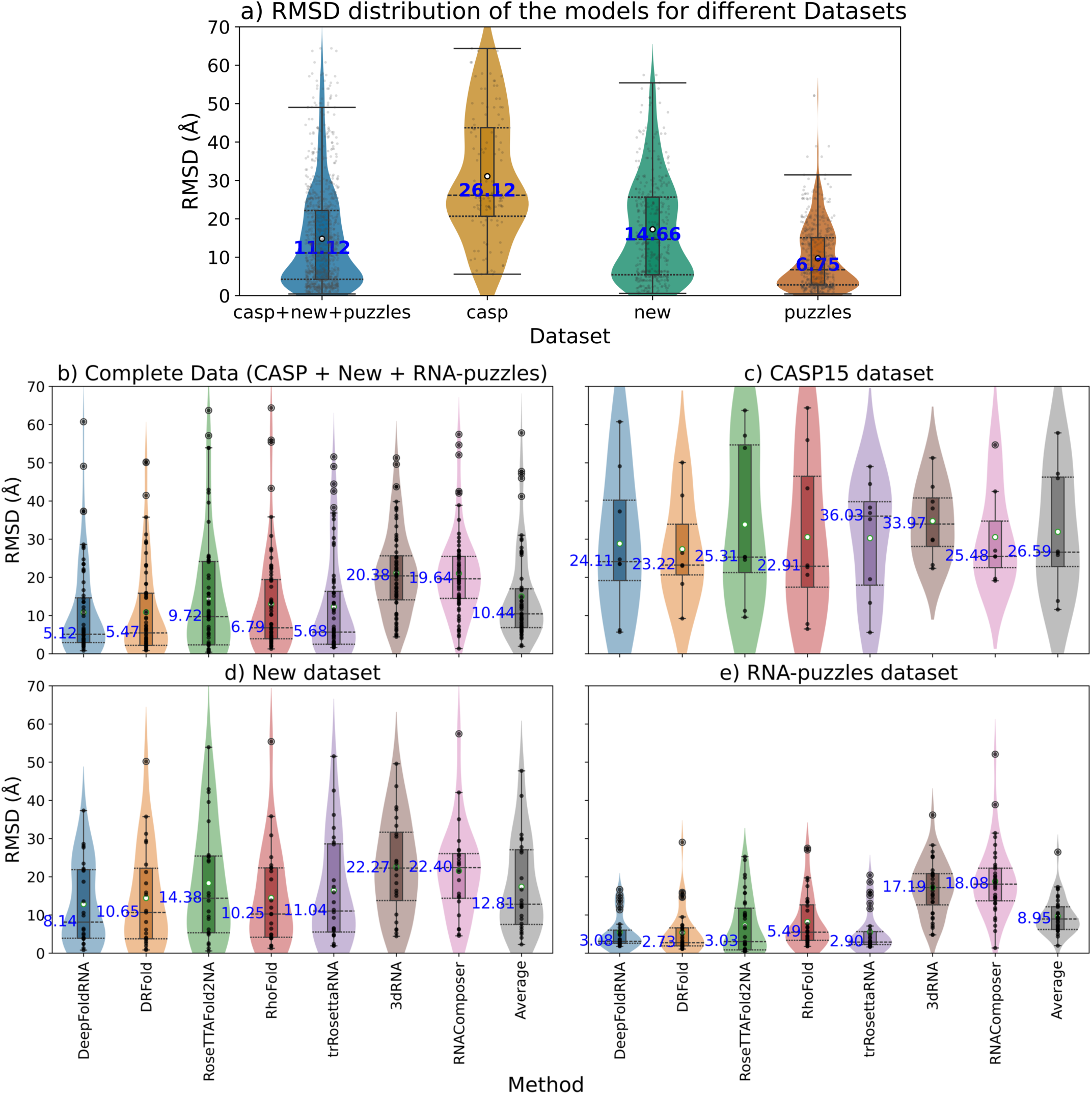
Box and violin plots showing the comparison of the methods based on different datasets. The median values are labelled with blue text and the whiskers denote the interquartile range **a)** The datasets are on the X-axis and the RMSD is on the Y-axis. We pooled all the models from different methods datasets together and only compared the the RMSD of the models based on their datasets. The median RMSD for the CASP dataset was the highest (26.12 Å), New dataset was in the middle (14.66 Å) and RNA-puzzles had the lowest median RMSD (6.75 Å). **b)** This plot shows the same comparison, but we look at each method separately. The Average plot (in grey) is the average of the models predicted by all the methods for a particular target. Generally, CASP15 dataset has the highest median RMSD for all methods, New dataset was in the middle and RNA-puzzles dataset has the lowest median RMSD for all the methods. The reason for RNA-puzzles being the easiest is because 35/36 targets are X-ray crstallographic structures and many of the targets were published before 2020, thus they might be included in the training sets of the ML-based methods. CASP dataset is the hardest because most of the targets are synthetic and Cryo-EM structures. The new dataset provides the most realistic performance estimates as it is a well-balanced dataset (comprising all kind of RNAs with representation from both X-ray crystallographic and Cryo-EM structures) and none of its targets are present in the training sets of the ML methods. DRFold has a median RMSD of 2.03 on RNA-puzzles dataset possibly because it has already seen most of the targets in the RNA-puzzles dataset while training thus giving an overinflated performance.

### Effect of RNA type on the prediction performance

In CASP15, most deep-learning based methods performed poorly on the synthetic targets in CASP15 and also on the RNA-protein complexes none of the methods were accurate. Therefore, we divided all the RNAs into three categories similar to the CASP15-RNA competition: Natural, Synthetic and RNA-protein complex and then compared the prediction performance according to RNA type. The median RMSD of all the models for natural targets was much lower than synthetic and RNA-protein complexes (fig 11a). We again looked at the same comparison but segregated the models based on data type. We observed that on CASP dataset, natural targets has the lowest median RMSD, RNA-protein targets were in the middle and synthetic targets had the highest median RMSD. For New dataset, natural has the lowest, while synthetic and RNA-protein complex had similar median RMSD (slightly higher for RNA-protein). Interestingly, on the RNA-puzzles this trend wasn’t observed (fig 11b) as the synthetic targets actually had lower median RMSD than natural and RNA-protein targets possibly because of the known problem with the RNA-puzzles dataset (old targets that are part of the training set of ML methods). We also looked at the performance dependence of the methods on the RNA type for all methods separately and found that for all methods (except DRFold) the natural targets are the easiest to predict, with RNA-protein being more difficult and synthetic being the hardest (fig 11c; S7).

**Fig 11.**
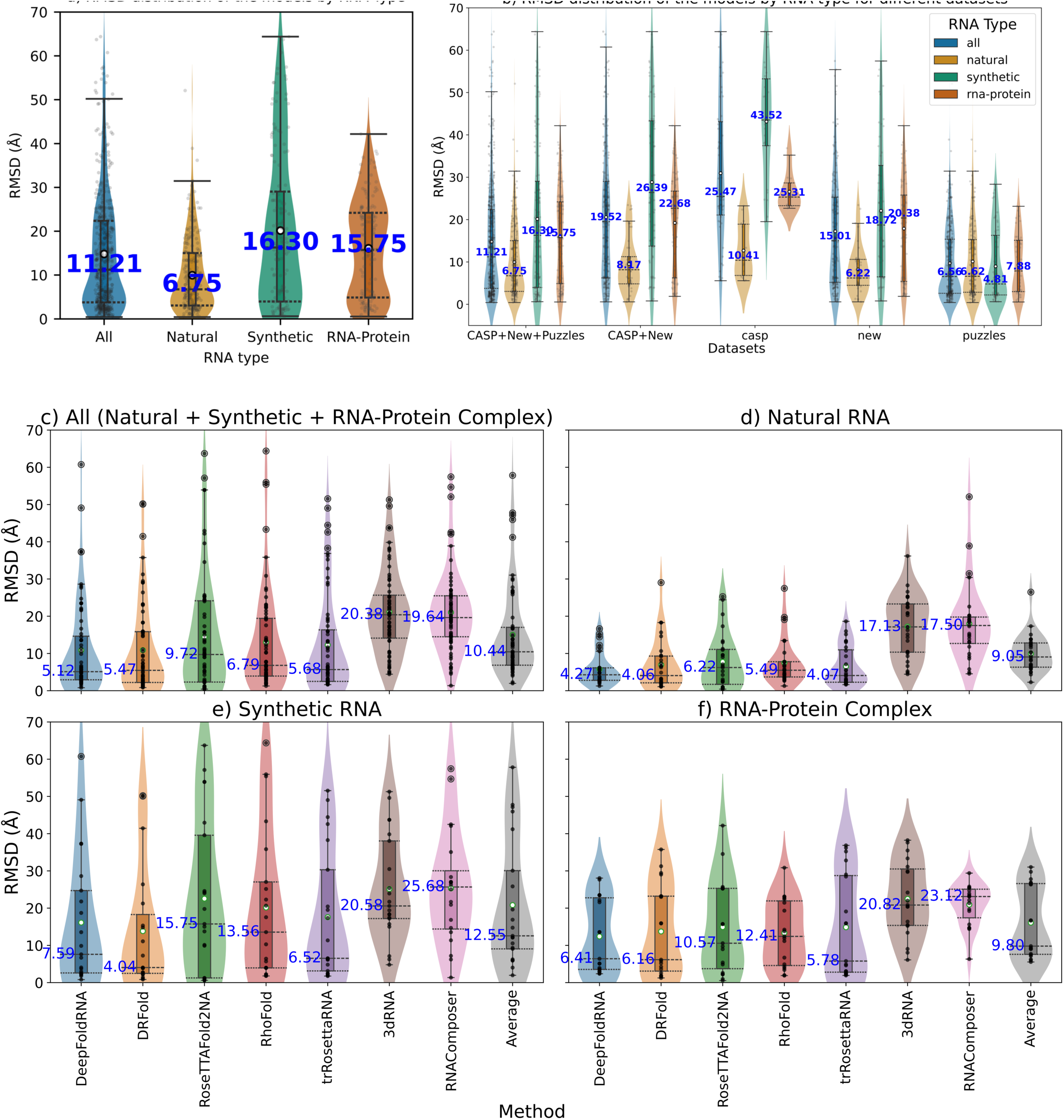
Comparing the prediction performance based on RNA type **a)** Violin plots showing the RMSDs of the models based on the RNA type. Natural targets have the lowest median RMSD (6.75 Å), RNA-protein have the second best (15.75 Å) and synthetic have the highest (16.30 Å). **b)** Performance difference for different RNA types based on different datasets. For CASP15 dataset, natural have the lowest, RNA-protein have the middle and the synthetic targets have the highest median RMSD. For the new dataset, natural have the lowest (6.22 Å), while RNA-protein and synthetic have similar median RMSDs with synthetic being slightly lower than RNA-protein (18.72 Å for synthetic and 20.38 for RNA-protein). Interestingly for the RNA-puzzles dataset the lowest median RMSD is for the synthetic targets (4.81 Å), while natural RNAs have slightly higher (6.62 Å) and the RNA-protein ones have the highest (7.88 Å). This discrepancy for this dataset is because many of the targets in this dataset are published pre-2020 so they are also present in the training sets of ML-methods thus resulting in an inflated performance (the difficulty of being a synthetic target doesn’t matter because ML-model has already learnt the structure). **c)** Performance comparison of all the methods separately for different RNA types. For all methods (except DRFold) the natural targets are the easiest to predict, with RNA-protein being more difficult and synthetic being the hardest based on the median RMSD scores.

### Dependency on the length of the RNA

We checked the correlation of the length of the target RNA with the RMSD of the predicted models for all the methods (fig 12a) and we did observe a positive correlation (as length increases, RMSD also increases). For longer RNAs, the RMSD of the predicted models was generally higher than that of the shorter RNAs. However, when we only looked at the correlation for shorter RNAS (RNAs with length < 100), the correlation was much weaker (fig 12b). On checking the correlation of RNAs of length > 100 with RMSD, we again observed a strong positive correlation (Fig S8). We observed a negative correlation between TMscore and the length of the RNA (as length increases, TMscore decreases) (fig 13a). In the previously published manuscripts [52], a positive correlation between TMscore and the length of the RNA (up to a certain length) has been observed. Therefore, we looked at the correlation between TMscore and length for RNAs of length less than 100 and RNAs of length more than 100 separately. We indeed saw a positive correlation between TMscore and RNA length for RNAs of length less than 100 (fig 13b). The correlation was again negative for RNA length > 100 (Fig S9). This indicates that both shorter and longer RNAs are difficult to predict because of certain confounding factors.

**Fig 12.**
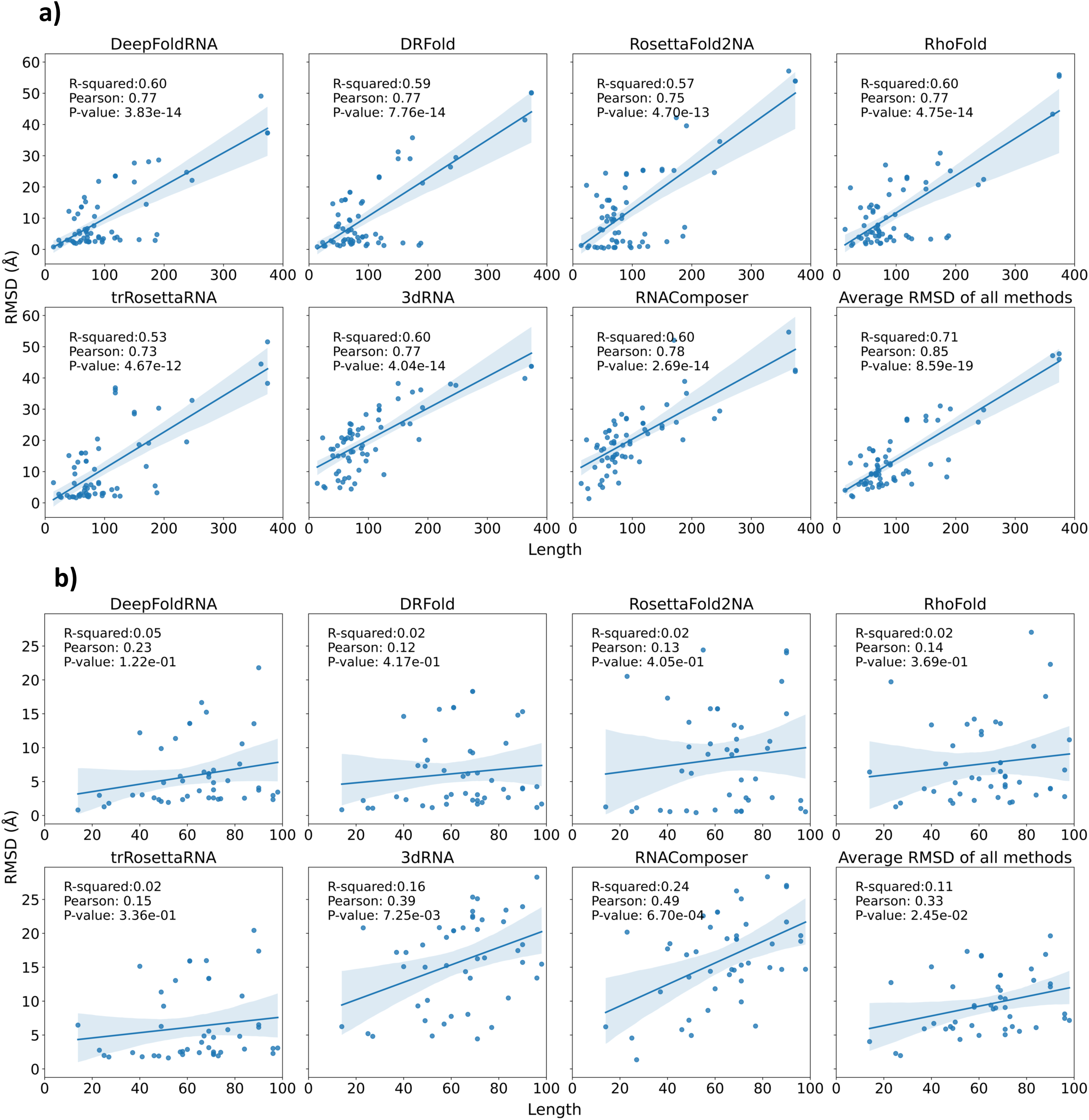
**a)** The correlation between the length of the target RNA and the RMSD of the predicted model for all the methods. In this plot RNAs of any length are considered. We observe a positive correlation between the length and the RMSD of the models for all the methods suggesting that longer the RNA, higher the RMSD of the predicted model and hence lower the quality of the predicted model. This shows that predicting the 3D structure of longer RNAs tends to be more challenging than that of shorter RNAs. **b)** The correlation between the length of the target RNAs and the RMSD of the predicted model for all the methods. In this plot, only RNAs with length < 100 are considered. We observe that the clear positive correlation that we observed in (a) for all models is only present for FA-based methods (3dRNA and RNAComposer) and is also much weaker than the first case. The correlation for ML-based methods is not there anymore. This suggests that on increasing the length of the target RNA (up to 100 nucleotides) there isn’t much effect on the quality of the predicted model.

**Fig 13.**
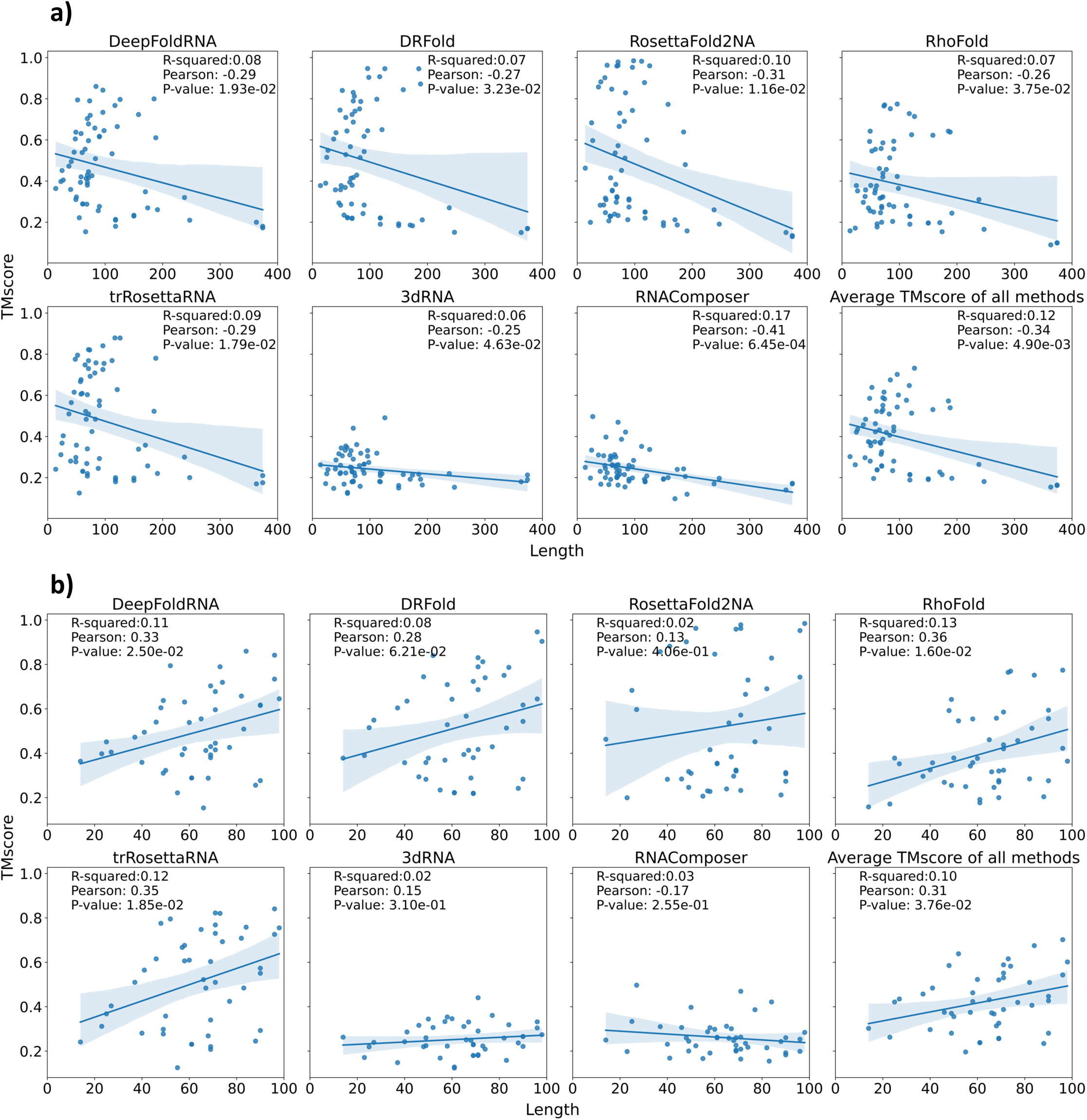
**a)** The correlation between the length of the target RNA and the TMscore of the predicted model for all the methods. In this plot RNAs of any length are considered. We observe a negative correlation between the length and the TMscore of the models for all the methods suggesting that longer the RNA, lower the TMscore of the predicted model and hence lower the quality of the predicted model. This could possibly be because because most ML-methods are trained on RNAs shorter than 200 nucleotides. **b)** The correlation between the length of the target RNAs and the TMscore of the predicted model for all the methods, in which only RNAs with length < 100 are considered. Contrary to what we expected, the weak negative correlation that we observed in (a) for all models is now a weak positive correlation for most methods. This suggests that on increasing the length of the target RNA (up to 100 nucleotides) the TMscore of the predicted model and hence the quality of the model also increases, which is unexpected, but has been previously reported by the RhoFold paper as well.

### Effect of the quality of MSA on the RNA

The depth of the MSA used as input to deep-learning-based structure prediction methods has previously been suggested to affect the quality of the predicted models [43]. Recently, Lee et al. found that expanding the MSA database boosts ColabFold’s CASP15 performance. [69]. Therefore, we investigated the effect of MSA quality on the RMSD of the predicted models.. The differences in the MSA tools (rMSA for DeepFoldRNA, rMSA-lite for RosettaFold2NA, rMSA + Infernal for trRosettaRNA, blastN for RhoFold) and the RNA databases that are used to create the MSA can cause a lot of variance in the quality of the MSA for the same target sequence [70,71]. We used the Log(N_eff_) measure to quantify the quality of the MSA depth. The N_eff_ metric is calculated from the hmmbuild command in the HMMAlign [72] tool and is a measure of the number of effective homologous sequences in the alignment. We don’t observe a correlation between the RMSD and the MSA depth for DeepFoldRNA and trRosettaRNA, but a negative correlation is observed for RoseTTAFold2NA and RhoFold, which indicates that higher-depth MSA is more informative and results in better quality prediction for these tools (fig 14a). If we look at the correlation between the TMscore of the targets and their MSA depth, we again observe a positive correlation for RoseTTAFold2NA and RhoFold, while that is not the case for DeepFoldRNA and trRosettaRNA (fig 14b).

**Fig 14.**
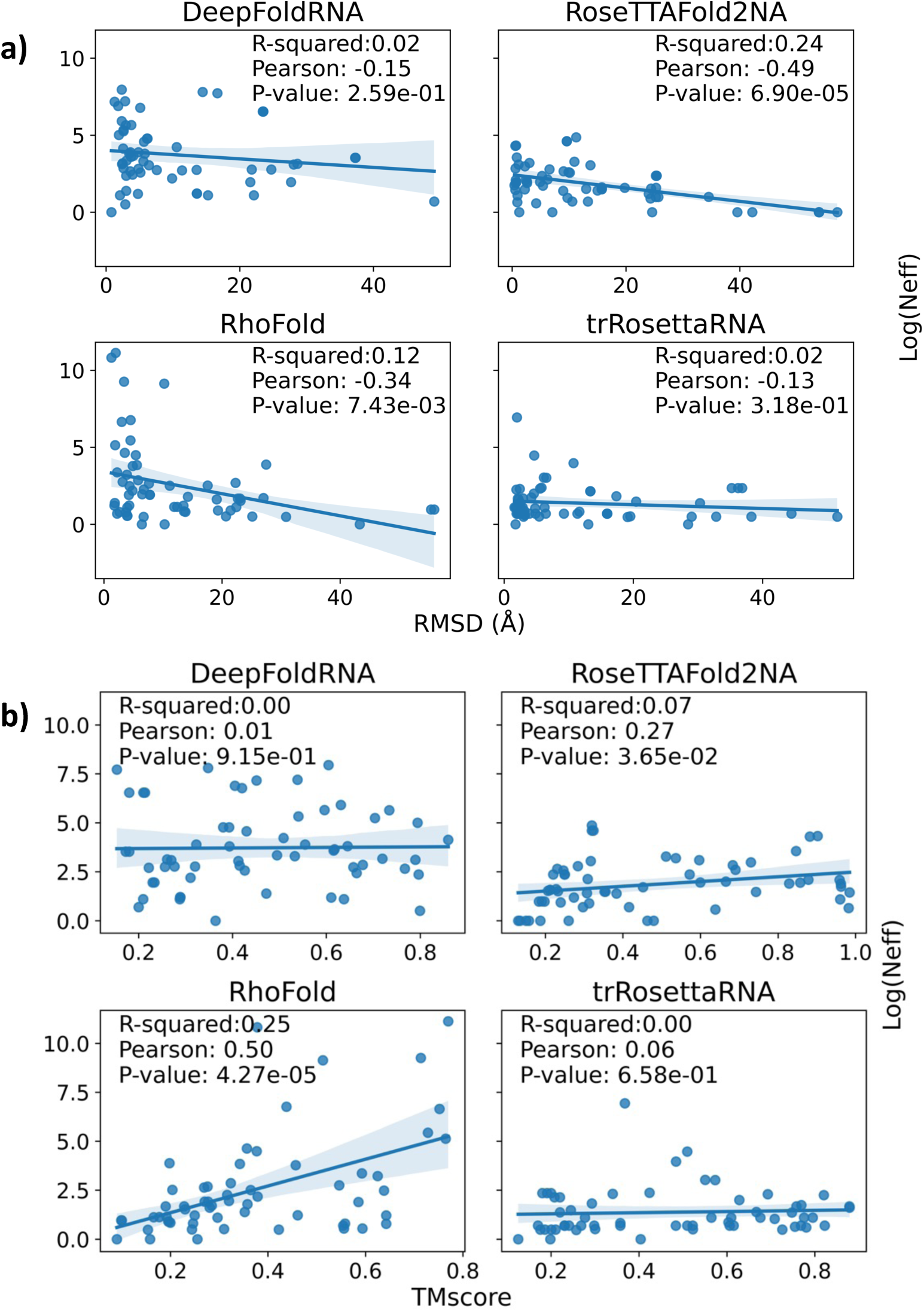
Scatter plots showing the correlation between RMSD/TMscore and the MSA depth (Log(N_eff_)) for the four ML-based methods (DeepFoldRNA, RosettaFold2NA, trRosettaRNA, RhoFold) that take MSA as input. DRfold was excluded as it doesn’t take MSA as input. As the methods used by the tools to create the MSA differ, we created separate scatter plots for each of the methods. DeepFoldRNA uses rMSA, RoseTTAFold2NA uses rMSA-lite, trRosettaRNA uses a mix of rMSA and Infernal, while RhoFold only relies on blastN to create the MSA. **a)** A negative correlation is observed for RoseTTAFold2NA and RhoFold which indicates that models of the targets with higher MSA depth have better quality (lower RMSDs). **b)** Scatter plots for the four methods between TMscore and the MSA depth (Log(Neff)). A positive correlation is observed for RoseTTAFold2NA and RhoFold which indicates that models of the targets with higher MSA depth have higher TMscore.

### Effect of secondary structure on the prediction performance

The secondary structure (ss) of the RNA plays an important role in the tertiary folding of the RNA molecule, therefore, it’s imperative to use the correct ss as input to the DL-model [73]. However, we also saw in the INF plots that the percentage of correctly predicted WC-pairs was on average already around 80% for the predicted models, which means that for most of the targets the correct ss information is already provided as input (predicted by respective third-party ss prediction tool). Nevertheless, we investigated how using the correct ss (extracted from the native PDB files) as input will improve the model prediction. We selected 14 targets that have their complete native structure available (no residues missing) and extracted the ss from their PDB files using the RNAPDBee 2.0 tool [74]. We observe that with the native ss as input the fragment-assembly-based methods (RNAComposer and 3dRNA) show a large improvement in the quality of the models, while for ML-based methods, we don’t see a huge difference (slight increase for DRFold and slight decrease in trRosettaRNA) in the final model quality (Table 8). The probable reason for this could be that the ML-based methods are actually trained on ss input from their respective ss prediction methods so ss extracted from native PDBs might not offer much improvement in the quality of the predicted model and native ss as input rather creates restraints that these tools are not able to handle, thus resulting in models with higher RMSds. We also tried removing the input ss information to the methods for one of the targets (8FZR) and predicted the final model without any input ss information. The quality of the final predicted model dropped substantially for all methods (except DRFold) on removing the ss information altogether (fig 15).

**Table 8.** RMSD comparison (in Å) of models for 13 targets predicted by methods that take secondary structure (ss) as input. The methods compared are RNAcomposer (rnac), 3dRNA (3dRNA), DRFold (drfold) and trRosettaRNA (trr). The columns without ‘_ss’ suffix indicate the predicted models where the input ss is the default one predicted by the respective associated ss prediction method for each tool (RNAfold for RNAComposer and 3RNA; SPOT-RNA for trRosettaRNA; PETfold + RNAfold for DRFold). The columns with ‘_ss’ suffix are the ones where the input ss is the one extracted from the native PDB file using the RNAPDBee tool. We observe that for both the fragment-assembly-based methods i.e. RNAComposer and 3dRNA, the average RMSD for predicted models is much lower with the native ss compared to the default ones (22.64 Å vs 13.72 Å for RNAComposer and 24.81 Å vs 20.65 Å for 3dRNA). For the ML-based methods i.e. DRFold and trRosettaRNA, we don’t see much difference between the two scenarios. We actually observe a slight increase in the average RMSD for DRFold (17.27 Å vs 18.19 Å) and a slight reduction in the average RMSD for trRosettaRNA (19.80 Å vs 18.63 Å). The probable reason for this could be that the ML-based methods are actually trained on ss input from their respective ss prediction methods so ss extracted from native PDBs might not offer much improvement in the quality of the predicted model and native ss as input rather creates restraints that these tools are not able to handle, thus resulting in models with higher RMSds.

| Target | rnac | rnac_ss | 3drna | 3drna_ss | drfold | drfold_ss | trr | trr_ss |
| --- | --- | --- | --- | --- | --- | --- | --- | --- |
| 7u0y | 15.45 | 12.02 | 8.1 | 9.26 | 6.03 | 7.98 | 4.18 | 5.02 |
| 7uri | 31.79 | 20.84 | 20.19 | 20.46 | 16.09 | 17.05 | 15.77 | 16.72 |
| 7wi9 | 16.25 | 16.09 | 17.19 | 16.39 | 16.4 | 16 | 17.19 | 17.18 |
| 7zj5 | 42.49 | NA | 44.18 | 38.76 | 50.2 | 53.47 | 51.98 | 39.23 |
| 8bgu | 23.9 | 14.32 | 24.95 | 10.22 | 3.57 | 3.46 | 4.53 | 4.37 |
| 8dp3 | 21.67 | 6.14 | 24.47 | 16.02 | 15.31 | 13 | 17.65 | 16.53 |
| 8eyu | 13.6 | 15.74 | 15.43 | 13.76 | 11.81 | 11.94 | 11.36 | 9.8 |
| 8fcs | 10.24 | 6.57 | 3.98 | 4.86 | 2.54 | 2.6 | 4.21 | 5.84 |
| 8gxb | 23.53 | 11.37 | 20.95 | 9.37 | 16.57 | 15.3 | 15.91 | 9.03 |
| r1107 | 19.78 | 12.51 | 22.77 | 22.82 | 18.52 | 17.41 | 13.24 | 6.67 |
| r1128 | 27.53 | 18.46 | 38.68 | 29.3 | 26.71 | 36.1 | 19.83 | 25.76 |
| r1138 | NA | NA | 51.33 | 48.07 | NA | NA | 44.44 | 45.15 |
| r1190 | 25.5 | 16.82 | 30.32 | 29.21 | 23.47 | 23.98 | 37.09 | 40.93 |
| **Average** | **22.64** | **13.72** | **24.81** | **20.65** | **17.27** | **18.19** | **19.8** | **18.63** |

**Fig 15.**
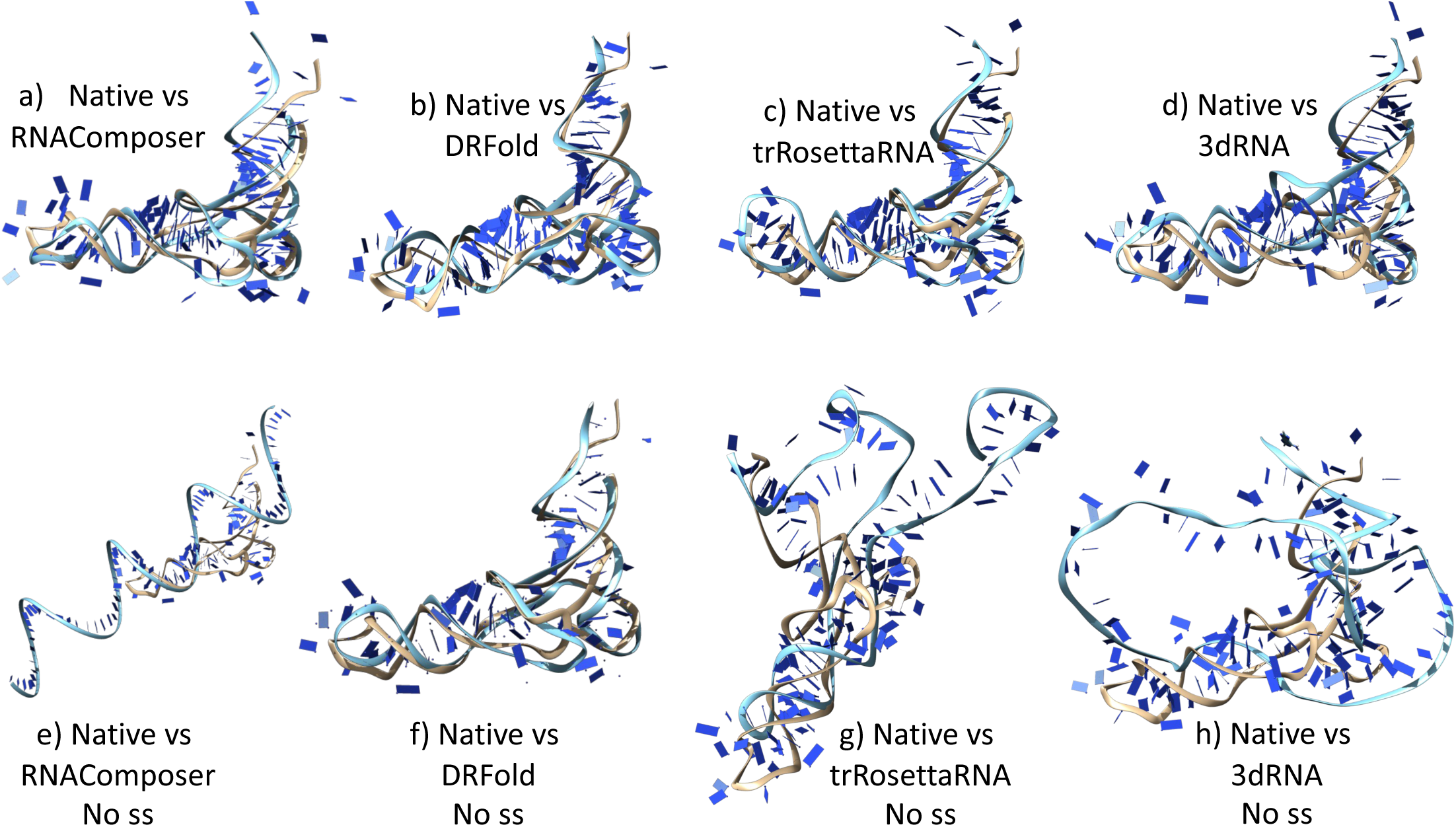
Superimposition of the native structure and predicted models of the target 8FZR showing the comparison between the quality of the predicted models by various tools when secondary structure (ss) is provided as input (predicted by the respective prediction method of each tool; SPOT-RNA for trRosettaRNA, RNAfold for RNAComposer and 3dRNA, and PETfold + RNAfold for DRFold) and when no secondary structure input is given. The native structure is shown in the light grey colour and the predicted models are in cyan. The first row of structures i.e. Fig a, b, c, and d show the superimposition from the case when secondary structure is provided as input and the bottom row i.e. Fig e, f, g, and h show the superimposition from the scenario when no secondary structure is provided as input. When ss is not provided as input, we can clearly see (in Fig e, f, g, h) that the quality of the predicted models by all the methods (except DRFold) is far worse than the models from the first row (in Fig a, b, c, d). This indicates that although replacing the predicted ss by extracted ss from native PDBs as input to these tools didn’t improve the quality of the final predicted model substantially, removing the ss as input altogether severely affects the quality of the final predicted model. Therefore, ss still plays a very important role in the accurate determination of the 3D RNA structure. The only method that wasn’t greatly affected by exclusion of ss as input was DRFold (possibly because it’s able to predict the nucleotide pairing and the associated restraints somewhat accurately even in the absence of ss because of how it’s trained the geometrical potentials it uses to fold the RNA; recall that it doesn’t take an MSA as input). The RMSD between the native and modelled structures are as follows: **a)** 5.70 Å for RNAComposer model **b)** 4.42 Å for DRFold model **c)** 5.78 Å for trRosettaRNA model **d)** 5.52 Å for 3dRNA model **e)** 59.46 Å for RNAComposer model without ss as input **f)** 4.63 Å for DRFold without ss as input **g)** 25.73 Å for trRosettaRNA model without ss as input **h)** 24.35 Å for 3dRNA model without ss as input

## Discussion and Conclusion

In this study, we conducted a comprehensive benchmarking of various RNA structure prediction methods across diverse datasets, each presenting varying levels of difficulty. While our primary focus was on deep-learning-based approaches, we also incorporated fragment-assembly methods to check the comparative effectiveness of machine learning (ML) versus traditional techniques. We evaluated seven RNA 3D structure prediction methods on three datasets, encompassing a total of 66 target RNAs. The benchmarked methods comprised five ML-based (deep-learning) approaches and two non-ML-based (fragment-assembly) methods. Performance assessment took into account multiple factors such as RNA length, MSA, dataset characteristics, and RNA types. Generally, ML methods exhibited significantly superior performance compared to their non-ML counterparts. The effectiveness of the methods varied based on the benchmarking dataset, with the CASP15 dataset posing the greatest challenge as this dataset included numerous targets that were either synthetic or RNA-protein complexes. In contrast, the RNA-puzzles dataset proved to be the easiest, possibly because it contains many targets published before 2020 which might have been part of the ML methods’ training datasets. Notably, the RNA-puzzles dataset predominantly comprised X-ray crystallography structures, while CASP15 mainly consisted of Cryo-EM structures. Given that X-ray crystallography structures generally have better resolution and accuracy, the ML methods, trained predominantly on crystallographic structures, exhibited better accuracy when modeling X-ray targets. The newly compiled dataset has a good balance of Cryo-EM and X-Ray structures where none of the targets were seen by the ML-methods previously and therefore this dataset provides the most realistic benchmarking set for evaluating the methods in a blind prediction scenario. On this dataset, DeepFoldRNA had the lowest median RMSD.

The performance was also affected by the type of RNAs and the length of the RNA. Natural RNAs were the easiest targets with best quality models, while RNA-protein complex models had medium accuracy, and the synthetic targets were the hardest. We also observed that generally the quality of the predicted models (based on RMSD) gets lower as the sequence length of the target RNA increases, although, the correlation is quite weak when looking at RNAs of length less than 100 nucleotides. Similar trend was seen when we looked at the correlation of length vs TMscore with longer RNAs having lower TMscore. However, we also observed a positive correlation between length and TMscore when only considering RNAs of sequence length less than 100. Generally, when trying to predict the structure of very long RNAs, the model quality is less accurate because the prediction tool is not able to predict long-range interactions, pseudo knots and the associated base-pairing. This is because getting a good quality MSA (non-sparse and high depth) for longer sequences is difficult as very few homologous sequence matches are found. However, when the sequences are too short then the deep-learning-models don’t have enough information to work with, diminishing the predicted model quality. This indicates that there is an optimum range for the length of RNA for the best prediction quality.

We also observed a slight correlation between the MSA and model quality for RosettaFold2NA and RhoFold methods, while there wasn’t any correlation observed for the other ML-methods (DeepFoldRNA and trRosettaRNA), which suggested that methods that only take MSA as input have worse performance than methods that take MSA + secondary structure (ss) as input. In simpler terms, the negative impact of using a lower quality MSA as input is more pronounced in the predictions generated by RoseTTAFold2NA and RhoFold compared to DeeFoldRNA and trRosettaRNA. One plausible explanation for this could be that DeepFoldRNA and trRosettaRNA utilize existing tools for predicting the secondary structure (PETFold for DeepFoldRNA and SPOT-RNA for trRosettaRNA) [75,76] and then use both the MSA and the secondary structure as input to their transformer network, while in the case of RoseTTAFold2NA and RhoFold the sole input to their neural network architecture is the MSA. Thus, even if the MSA is sparse, DeepFoldRNA and trRosettaRNA can compensate by leveraging the input ss information to predict the correct 1D and 2D geometries, whereas RoseTTAFold2NA and RhoFold have to rely solely on the MSA, which might lead to comparatively higher inaccuracies in predicting the base-pairing information and the associated restraints. These inaccuracies propagate downstream into the final prediction resulting in models with lower TMscore.

We also checked the dependence of the model quality on the input ss. Although, using ss extracted from native PDBs instead of predicted ss as input didn’t improve the predicted model quality for the ML-based methods by much, removing the input ss altogether reduced the model quality substantially for most methods (except DRFold). This underscores the importance of incorporating ss information in RNA structure prediction, even if the predicted ss may not always enhance model quality substantially.

Altogether, the DL-based methods had the best predicted models with DeepFoldRNA being the best followed by DRFold as the close second. The DL-based methods had the best predicted models for most targets except for the synthetic targets in CASP15 dataset. The FA-based methods lagged behind for most targets and we recommend to rely on the ML-based methods for predicting RNA 3D structures moving forward. When working with synthetic targets, all of the methods have a poor performance and none of them can predict a reliable 3D structure. For future method development, we recommend to develop methods that use both MSA and secondary structure as input. Increasing the MSA depth by using metagenomic datasets has shown improvements in case of AlphaFold [69] and we recommend to follow suit for RNA structure prediction as well. Deeper MSAs for RNAs can be constructed using the new RNA database published by Chen at al. [77]. To improve the input ss we recommend to use a consensus ss predicted by multiple methods, especially the latest ML-based methods [78]. Even the best median INF-nwc score among all the methods was less than 50%, which suggested that none of the methods are able to predict the non-Watson-crick pairs accurately. Therefore, including ss prediction methods that can predict non-canonical base pairs, pseudoknots and other complex RNS ss elements more accurately should also be taken into account for future method development. Last but not the least, as more RNA experimental structures are solved and the size of the RNA PDB database increases with better representation of RNAs from previously uncharacterized Rfam families, the ML-based methods will improve as well.

## Data availability

The FASTA sequences of the target RNAs from all the datasets, the PDB files of the predicted models by all seven methods, different comparison metrics and the comparative analysis scripts are all available at https://github.com/akashbahai/rna_benchmarking. The benchmarked prediction methods are available at the following links:

RoseTTAFold2NA at https://github.com/uw-ipd/RoseTTAFold2NA DeepFoldRNA at https://github.com/robpearc/DeepFoldRNA DRfold at https://github.com/leeyang/DRfold/

RhoFold at https://github.com/Dharmogata/RhoFold trRosettaRNA at https://yanglab.qd.sdu.edu.cn/trRosettaRNA/ RNAComposer at https://rnacomposer.cs.put.poznan.pl/ 3dRNA at http://biophy.hust.edu.cn/new/3dRNA

## Acknowledgements

The computational work for this article was performed on resources of the National Supercomputing Centre (NSCC), Singapore (https://www.nscc.sg).

## Key Points

- Systematic benchmarking of five latest deep-learning and two fragment-assembly based methods on diverse datasets
- Compiled a new balanced dataset with latest RNA structures for benchmarking
- Generally, the ML-based methods outperform the traditional fragment-assembly based methods with DeepFoldRNA having the best predicted models overall
- The performance of the methods is dependent on the MSA depth, RNA type, and secondary structure.

