## Supplementary for "Systematic benchmarking of deep-learning methods for tertiary RNA structure prediction"

**S1. RNA Structure Prediction Methods**

**S1.1 RosettaFold2NA:** RoseTTAFold2NA [12] is a deep learning-based method for predicting the 3D structure of RNA and protein-nucleic acid complexes. It is an extension of the original RoseTTAFold [13,14] method, which was developed for protein structure prediction. RoseTTAFold2NA uses a single trained network to produce 3D structure models with confidence estimates for protein-DNA and protein-RNA complexes, and for RNA tertiary structures. The architecture of RoseTTAFold2NA is based on a deep neural network that combines convolutional and recurrent layers to capture both local and global features of the input sequences. The input to the network consists of multiple sequence alignments (MSAs) of related protein and nucleic acid molecules, which are generated using sequence similarity searches against RNA sequence databases using rMSA lite.

**S1.2 DeepFoldRNA:** DeepFoldRNA is a fully automated method for predicting RNA tertiary structures from sequence alone, using deep self-attention neural networks to predict geometric restraints. It takes the MSA and predicted secondary structure as input and then uses a self-attention-based neural network architecture to predict geometric restraints, which are then used to guide the construction of 3D RNA structures through limited-memory Broyden-Fletcher-Goldfarb-Shanno (L-BFGS) minimization simulations. The geometric potential is derived from the predicted distances between pairs of atoms in the RNA molecule. Specifically, the predicted geometric restraints include pairwise distance maps between the N1/N9 atoms, C4’ atoms, and backbone P atoms, as well as inter-residue and backbone torsion angles (𝜔, 𝜆, 𝜂, 𝜃). These predicted geometric restraints are converted into composite potentials by taking the negative log-likelihood of the binned probability predictions, which are then used to guide the L-BFGS folding simulations.

**S1.3 DRFold:** DRfold is a novel deep learning-based method for ab initio RNA structure prediction that uses self-attention transformer networks to learn coarse-grained RNA structures directly from sequence. DRfold adopts a coarse-grained model of RNA specified by the phosphate P, ribose C4’, and glycosidic N atom of the nucleobase for training efficiency. The predicted conformations are further optimized by a separately trained deep-geometric potential through gradient-descent based simulations.

**S1.4 RhoFold:** RhoFold is an end-to-end deep learning method for accurate de novo RNA 3D structure prediction. It utilizes multi-aspect information of the RNA sequence to infer the 3D structure, including multiple sequence alignment (MSA) and information from a newly proposed RNA foundation model (RNA-FM). The method also introduces secondary structure information into the loss function and employs a novel procedure to perform self-distillation. The pipeline consists of three main components: the feature extraction module, the structure prediction module, and the structure refinement module. The entire model is fully differentiable, and several additional constraints are added to the structure output to ensure the final output is valid without clashing structures. The feature extraction module uses an RNA foundation model (RNA-FM) and a 4-layer E2Eformer module to learn the sequence representations and interactions between different nucleotides. The structure prediction module uses an 8-layer structure module to generate the final RNA 3D structures. Finally, the structure refinement module employs a recycling technique similar to AlphaFold to enhance prediction accuracy.

**S1.5 trRosettaRNA:** This method is built on top of trRosetta method (for protein structure prediction) and is adapted for RNAs. The authors used rMSA to build a MSA and SPOT-RNA method for secondary structure prediction. The deep-learning architecture of the method is a transformer network (named RNA-former) that takes MSA and pair representation as input and predicts 1D and 2D geometries, which are then converted into restraints to guide the 3D structure folding step. The full-atom structure of the RNA is generated by energy minimization with deep learning potentials and physics-based energy terms from Rosetta. The authors also constructed a self-distillation dataset by collecting the bpRNA sequences from the Rfam database.

**S1.6 RNAComposer:** This is a fragment-assembly based method that takes the RNA sequence and secondary structure as input to predict the RNA 3D structure. It works by breaking the secondary structure of the RNA into multiple substructures, including helices, loops, and bulges. It then searches for matches for each of the substructures in a database of known helices and loops from experimentally determined 3D structures, called RNA FRABASE. The matches are selected based on the secondary structure topology, sequence similarity and source structure resolution. The 3D structure is then assembled by superimposing common Watson-crick pairs at the ends of the substructures to combine all the substructures together and get the full-length 3D structure. The assembled structure is refined and energy minimized to obtain the final predicted model. The webserver for this method is available at <http://rnacomposer.ibch.poznan.pl/>. The webserver provides multiple choices for the method to be used to predict the secondary structure. However, we used RNAFold method to predict the input secondary structure for our benchmarking.

**S1.7 3DRNA:** This method also works similar to RNAComposer i.e. using secondary structure elements as building blocks to assemble the complete 3D structure of the RNA. The secondary structure is broken down into substructures of helices, different loops, and pseudoknots. These substructures are searched in a library of 3D structure-based templates assembled from known experimental RNA structures. In case of no matches, the templates are built using Bi-residue or distance geometry algorithm. The matched templates are then assembled to create initial 3D structures, which are put through a simulated annealing Monte Carlo process. This involves iteratively translating or rotating randomly selected movable elements, incorporating experimental restraints when available, and clustering the resulting structures using k-means clustering. The centroids of the clusters can be scored using a statistical potential to rank the structures and select the best predictions. The method is available at <http://biophy.hust.edu.cn/new/3dRNA>. The method can use multiple methods to predict the secondary structure, and we used RNAFold again to predict the secondary structure in this benchmarking.

**S2. CASP 15 results**

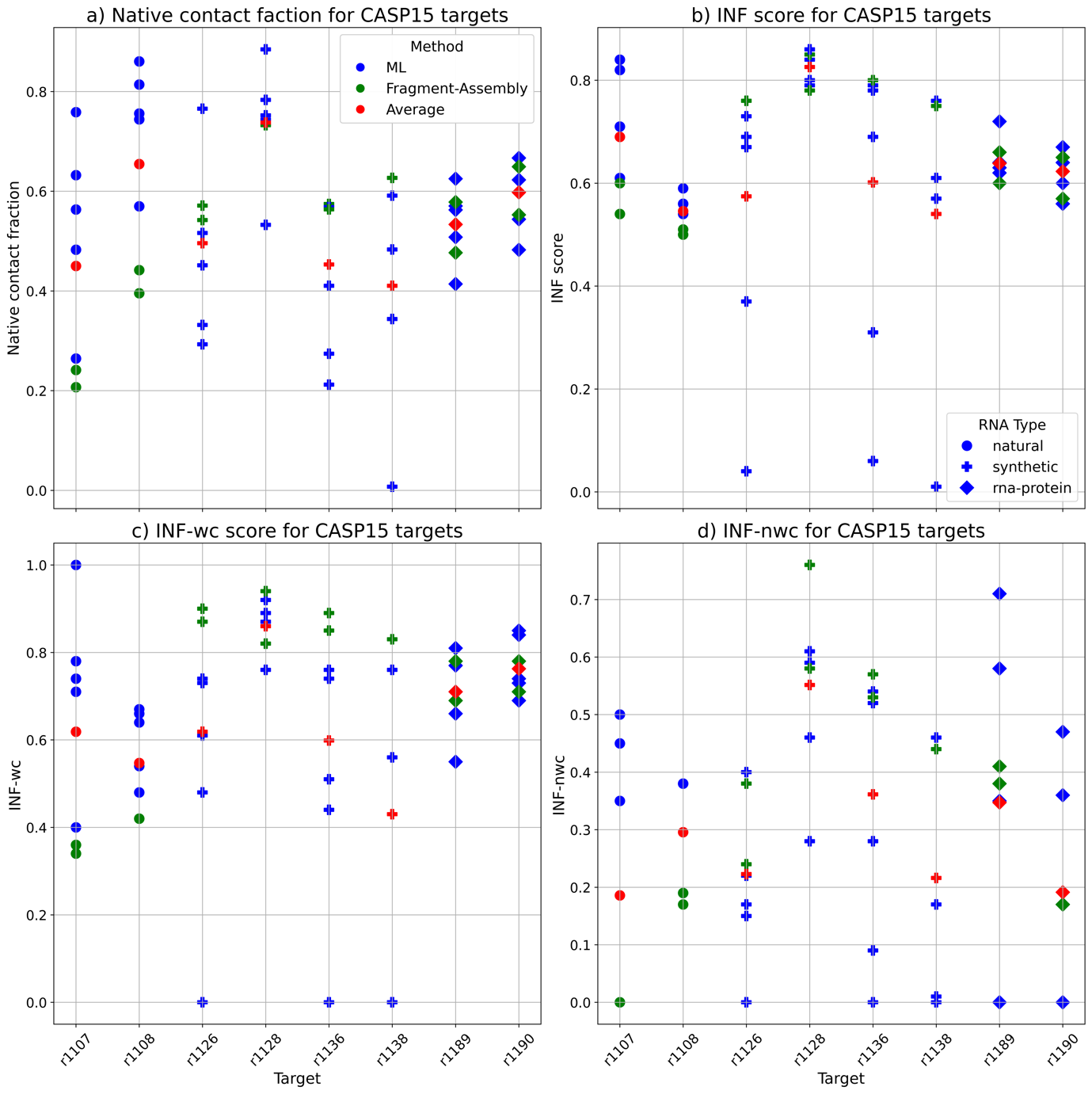
 **Fig S2.** Scatterplot showing the performance of the predicted models based on various metrics for the targets in the CASP15 dataset.

**S3. New dataset results**

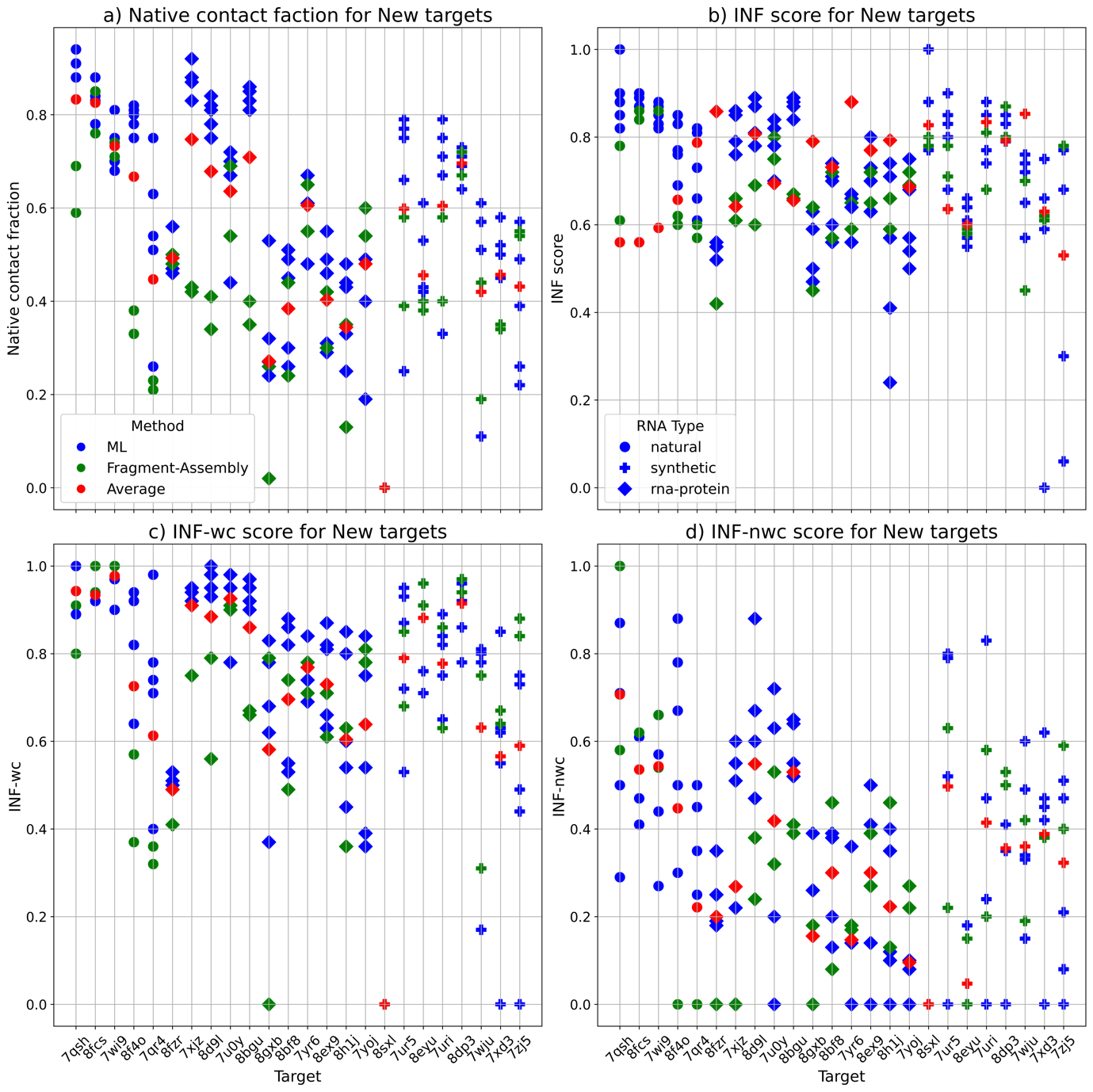
 **Fig S3.** Scatterplot showing the performance of the predicted models based on various metrics for the targets in the New dataset.

**S4. RNA-puzzles dataset results**

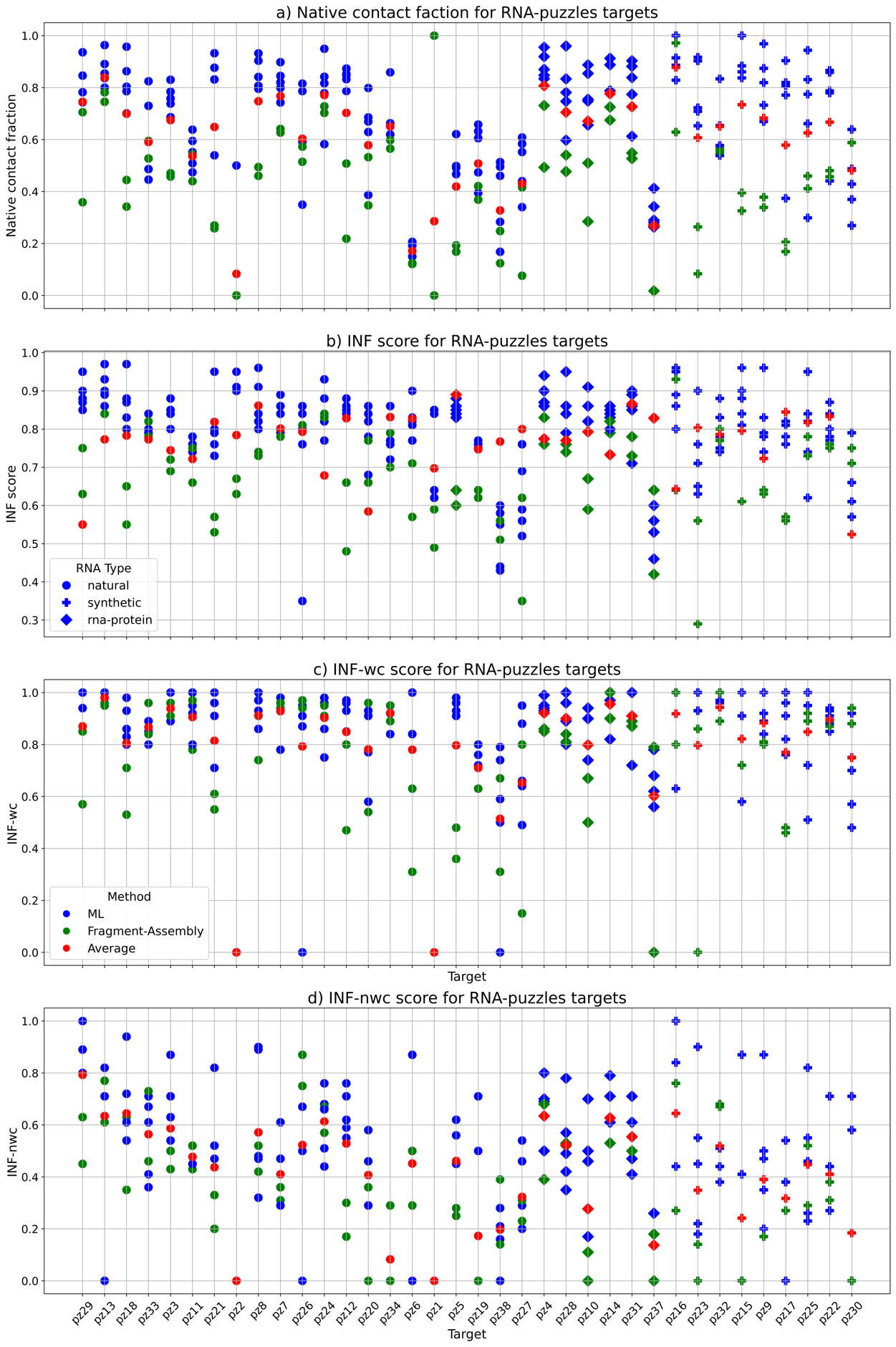

**Fig S4.** Scatterplot showing the performance of the predicted models based on various metrics for the targets in the RNA-puzzles dataset.

**S5. Pairwise comparison of different methods based on TMscore**

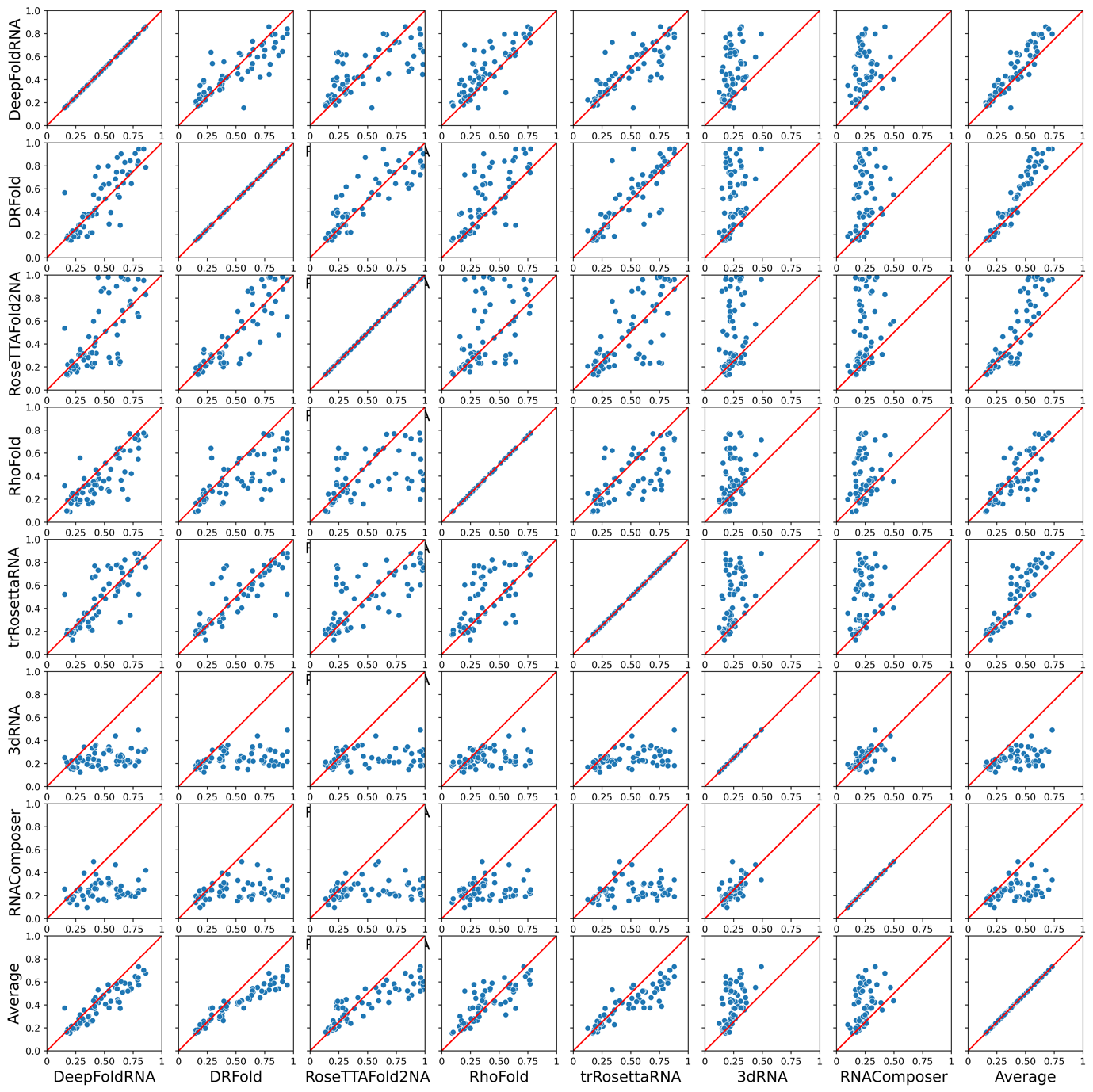

**Fig S5.** Scatterplot showing the performance comparison of each method against every other method on all the targets based on TMscore. If a point lies on the red-coloured x=y line, it indicates that the TMscore of the predicted model from both the methods is exactly the same i.e. they have similar prediction performance for that target. Points above that line indicate a higher TMscore for the model predicted by the method on the y-axis (i.e. method on the y-axis is better) and points below that line indicate vice-versa. Most of the ML-based methods have a better performance than the average prediction (last row of plots), while the FA-based methods are much worse than the average prediction (Average vs 3dRNA and Average vs RNAComposer plots in the last row). When compared against all other methods using the TMscores of the predicted models, DeepFoldRNA and DRFold are the best methods (DeepFoldRNA is slightly better).

**S6. RMSD cut-off plot for all the methods based on different datasets**

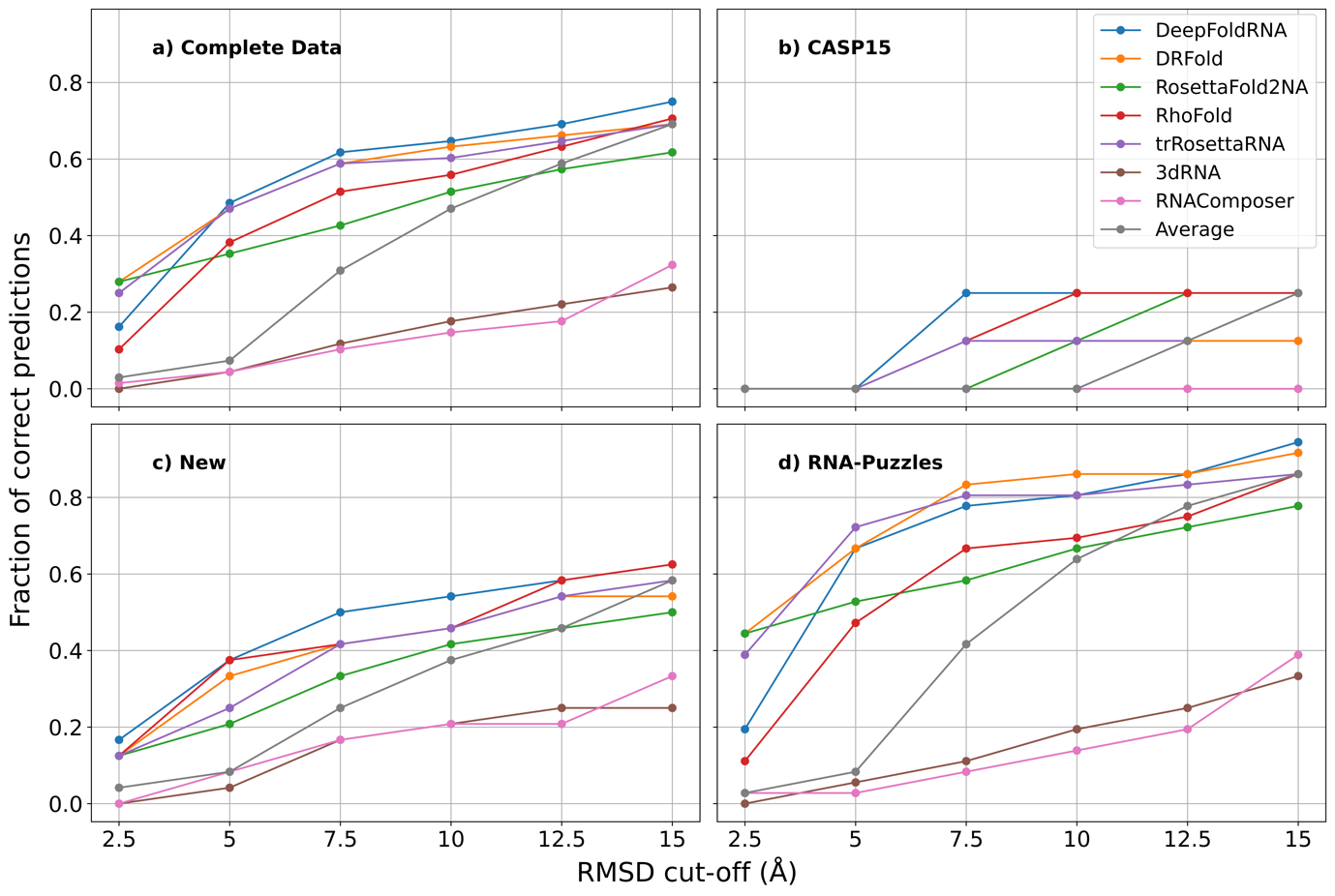
 **Fig S6.** RMSD cut-off plots for the seven methods based on the datasets they are benchmarked on. At an RMSD cut-off of 10 Å, in CASP15 none of the methods are able to even predict 30% of the targets correctly, while in the New dataset most ML-methods are able to correctly predict about 40% or more targets correctly (DeepFoldRNA predicts almost 50% targets correctly, the Average (in grey) correctly predicts about 38% targets correctly). In the RNA-puzzles dataset, most ML-methods have a correct prediction rate of 60% or higher with some even surpassing 80% (DRFold, DeepFoldRNA and trRosettaRNA). The Average method (in grey) for RNA-puzzles dataset at 10 Å predicts 62% of the targets correctly. This overinflated performance of ML-methods for RNA-puzzles dataset is because they have many of the targets in their training set.

**S7. RMSD cut-off plot for all the methods based on different RNA types**

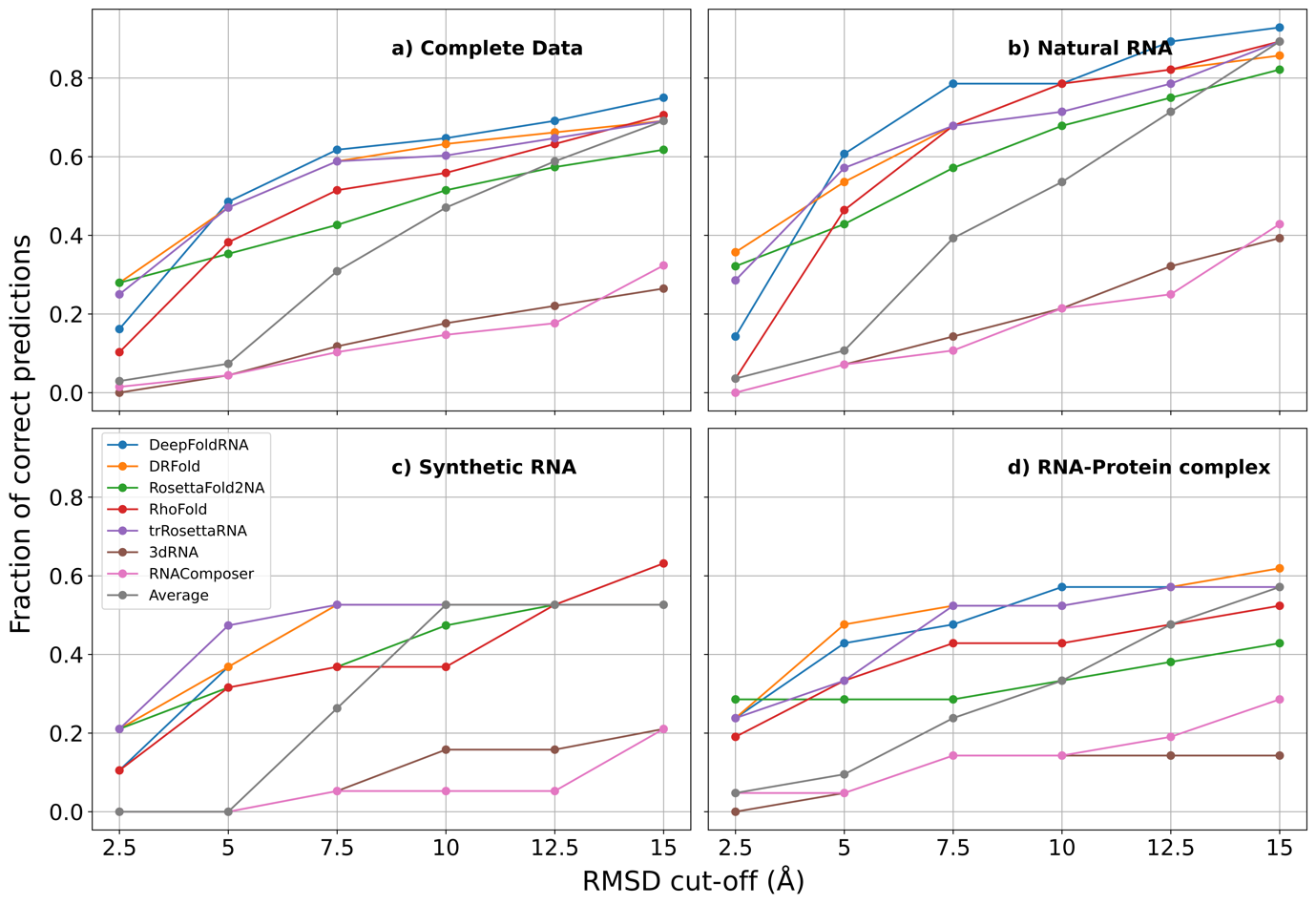

**Fig S7.** RMSD cut-off plots for the seven methods based on the RNA type. At an RMSD cut-off of 10 Å, for Natural RNAs the ML methods are able to predict 65% to 80% of the targets correctly (DeepFoldRNA and RhoFold are able to predict almost 80% of the natural RNA targets correctly), while in the case of Synthetic and RNA-protein complexes the % of correctly predicted targets is much lower (50% for synthetic by the best method and 59% for RNA-protein complex).

**S8. Correlation between RMSD and Length of the RNAs for RNAs longer than 100 nucleotides**

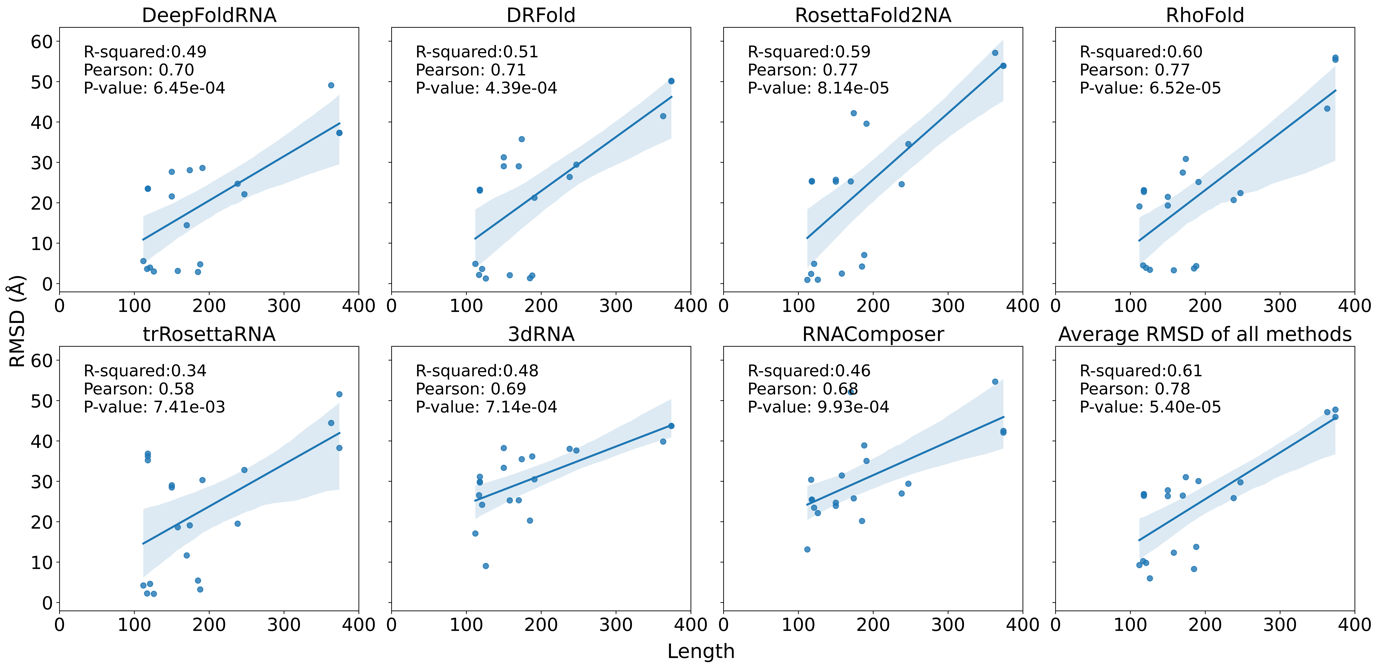

**Fig S8.** The correlation between the length of the target RNAs and the RMSD of the predicted model for all the methods. In this plot, only RNAs with length > 100 are considered. We see a positive correlation between RMSD and length indicating that as the RNA length increases the model quality decreases.

**S9. Correlation between TMscore and Length of the RNAs for RNAs longer than 100 nucleotides**

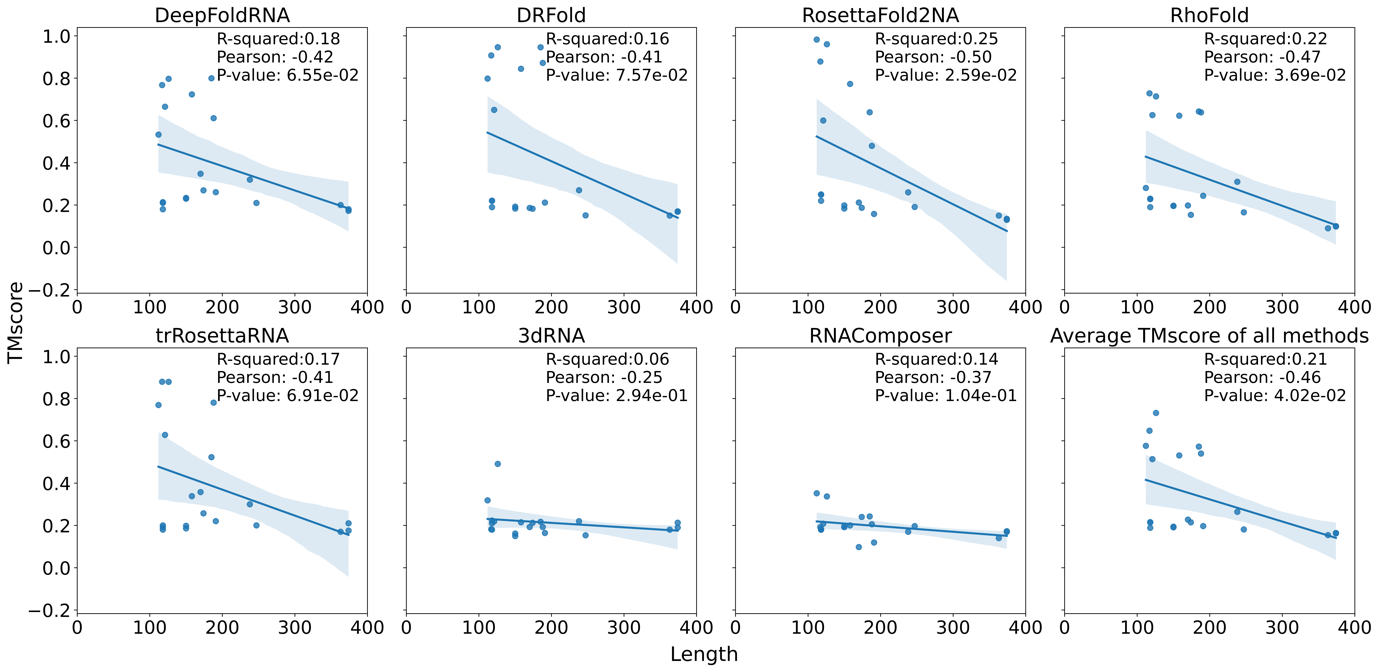

**Fig S9.** The correlation between the length of the target RNAs and the TMscore of the predicted model for all the methods. In this plot, only RNAs with length > 100 are considered. We see a negative correlation between RMSD and length indicating that as the RNA length increases the TMscore decreases, thus the model quality also decreases.
